# Quantifying and cataloguing unknown sequences within human microbiomes

**DOI:** 10.1101/2021.01.22.427751

**Authors:** Sejal Modha, David L. Robertson, Joseph Hughes, Richard J. Orton

**Author notes:** Corresponding author: Sejal Modha. Indicates last author (Joseph Hughes, Richard J. Orton).

## Abstract

Advances in genome sequencing technologies and lower costs have enabled the exploration of a multitude of known and novel environments and microbiomes. This has led to an exponential growth in the raw sequence data that is deposited in online repositories. Metagenomic and metatranscriptomic data sets are typically analysed with regards to a specific biological question. However, it is widely acknowledged that these data sets are comprised of a proportion of sequences that bear no similarity to any currently known biological sequence, and this so-called ‘dark matter’ is often excluded from downstream analyses. In this study, a systematic framework was developed to assemble, identify, and measure the proportion of unknown sequences present in distinct human microbiomes. This framework was applied to forty distinct studies, comprising 963 samples, and covering ten different human microbiomes including fecal, oral, lung, skin and circulatory system microbiomes. The framework was used to determine the proportion of taxonomically unknown sequences present within samples, and to compare such sequences both within and across assembled metagenomes. We found that whilst the human microbiome is one of the most extensively studied, on average 2% of assembled sequences have not yet been taxonomically defined. However, this proportion varied extensively among different microbiomes and was as high as 25% for skin and oral microbiomes that have more interactions with the environment. The publicly available data sets used have not previously been systematically mined to quantify and compare such dark matter. Typically, these unknown sequences are found in several microbiomes and potentially belong to unidentified novel microbes that we interact with on a daily basis. A cross-study comparison led to the identification of similar unknown sequences in different samples and/or microbiomes. A rate of taxonomic characterisation of 1.64% of unknown sequences being characterised per month was calculated from these taxonomically unknown sequences discovered in this study. Additionally, the approach led to the discovery of several potentially novel viral genomes that bear no similarity to sequences in the public databases. Both our computational framework and the novel unknown sequences produced are publicly available for future cross-referencing.

## Introduction

Metagenomics has become an increasingly mainstream tool to catalogue the microbial makeup of any given habitat (Aguiar-Pulido et al., 2016; Koonin, 2018; Quince et al., 2017; Thomas and Segata, 2019). It has been applied to a diverse range of environments from human body sites (Foulongne et al., 2012; Gevers et al., 2012; Consortium et al., 2012; Qin et al., 2010) to the depths of vast oceans (Breitbart et al., 2002; Hurwitz and Sullivan, 2013; Mizuno et al., 2013). Metagenomics, compared to culture based methods, provides a relatively unbiased approach to observe, measure and understand the interactions of the microbes within communities as well as with their hosts (Quince et al., 2017). Underpinned by relatively cheap sequencing costs and providing powerful insights, metagenomic has become a routine technique to study the microbial content of any environment (Koonin, 2018).

These advances in sequencing technologies and the importance of data sharing for reproducible research have led to the rapid expansion of publicly available sequence data. This has led to a rapid growth in online sequence databases such as GenBank, that store nucleotide and protein sequence data from various organisms (Cochrane et al., 2015; Karsch-Mizrachi et al., 2017). However, although the raw sequences generated as part of metagenomic experiments are made publicly available through the Short Read Archive (SRA) or European Nucleotide Archive (ENA) repositories, the complete set of assembled contigs from a study are rarely submitted to online databases (Connor et al., 2019). The reason for the absence of this type of data can be associated with the shear number of contigs generated and the requirement for sequences to be annotated before their submission, which is difficult when the organism the sequence came from is unknown, and when the number of contigs is large. Additionally, taxonomically unidentifiable contigs are typically discarded and excluded from downstream analyses (Figure 1(a)), but such sequences represent novel, and potentially widespread lifeforms and cataloguing their sequences and where they are found will aid taxonomic classification and our understanding of their biological nature in the future.

**Figure 1:**
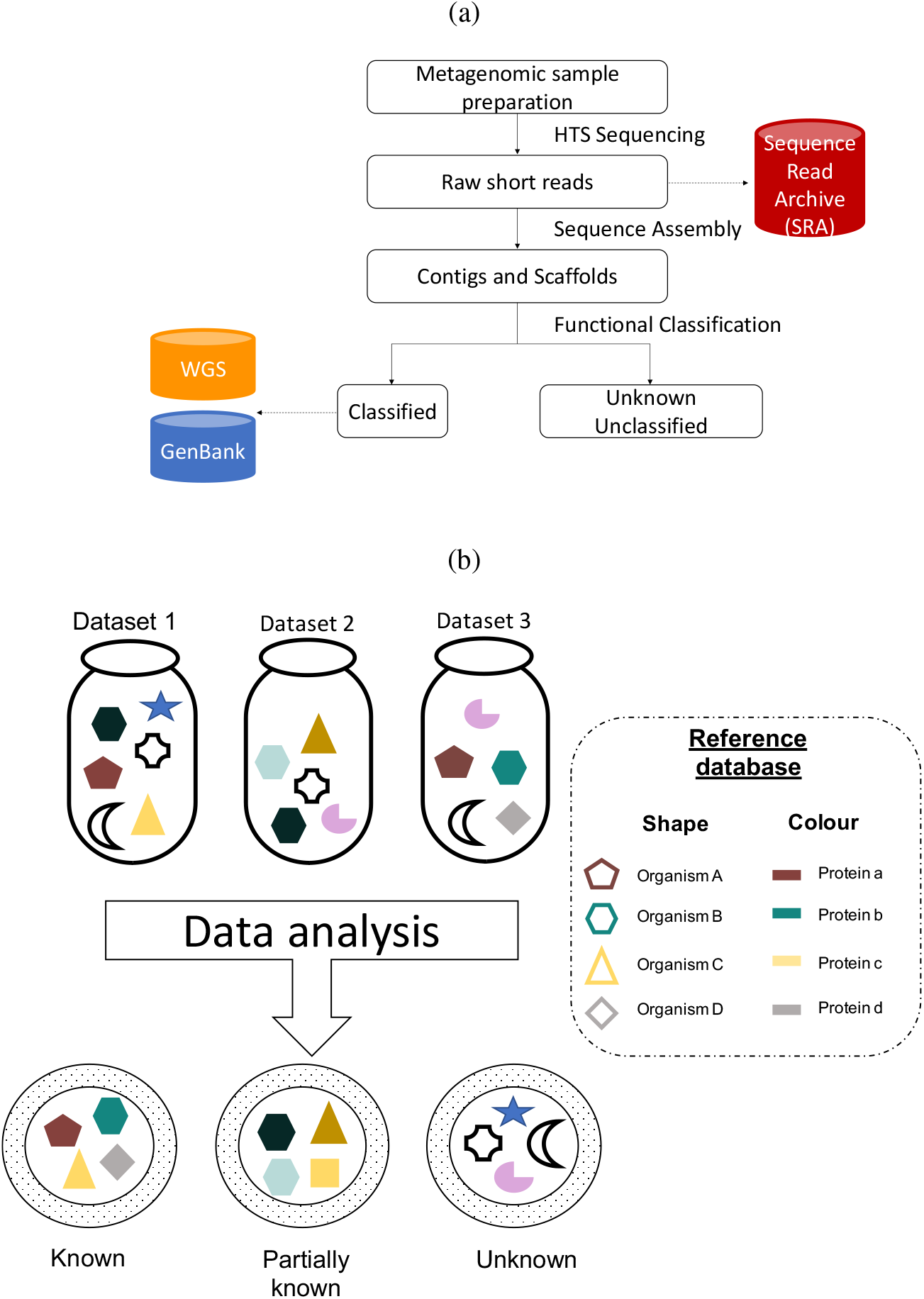
Overview of existing metagenomic analytical workflow and the definition of unknown sequence matter. (a) Typical metagenomic analytical workflow with data submission steps. (b) A schematic representation of known, partially known and unknown sequence matter in the metagenomic data sets.

**Figure 2:**
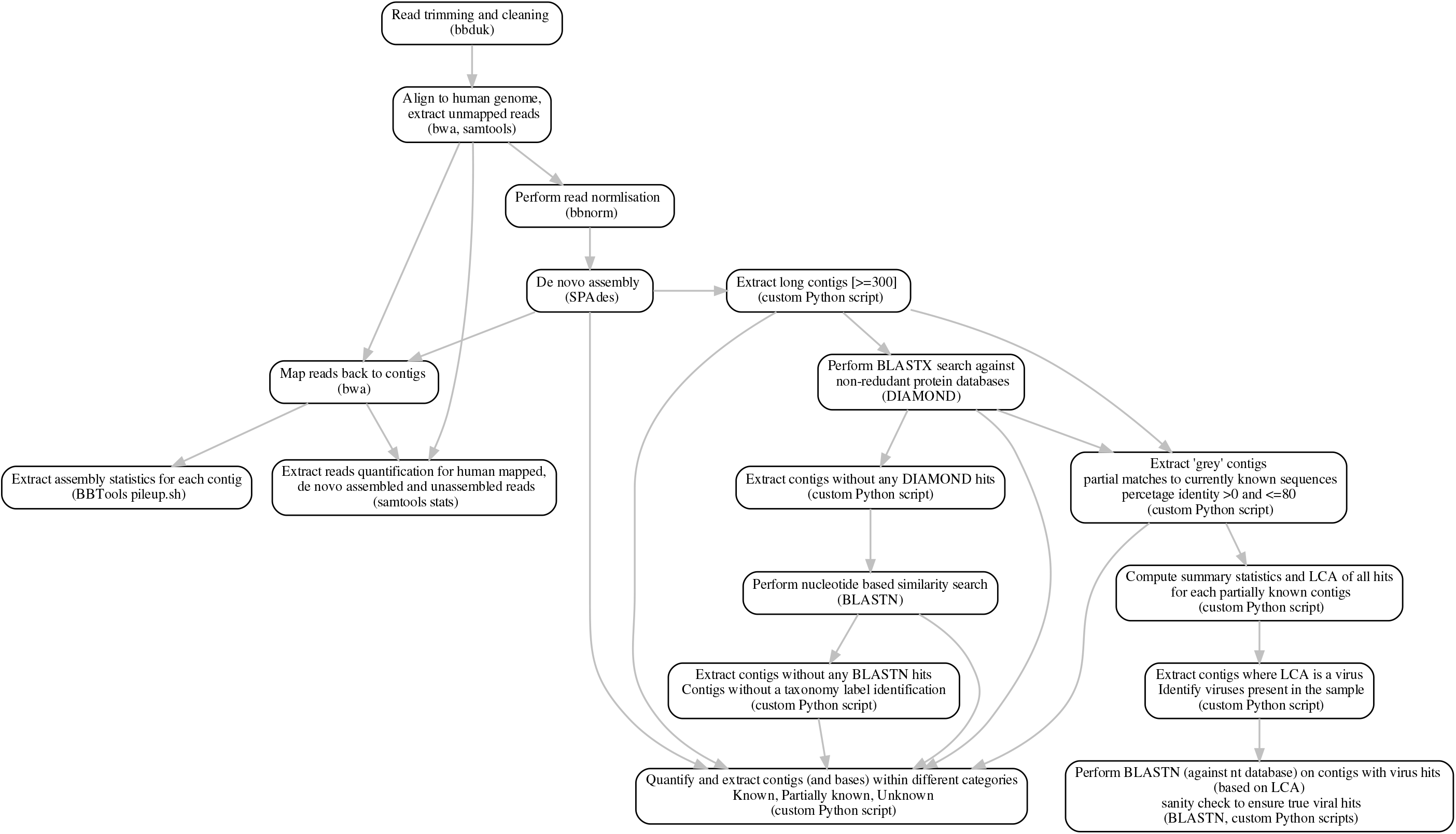
Detailed workflow of the metagenomic analysis and unknown sequence identification pipeline.

The raw data in public databases is typically analysed using metagenomic protocols designed to address specific biological questions. There are a range of different tools and pipelines available for metagenomic sequence analysis, but there are limited comparisons of these pipelines as they are usually developed to address specific research questions. For example, there are approximately 50 workflows available for virus metagenomic analysis that were used in different publications with primarily different aims (Nooij et al., 2018). As part of a routine metagenomic analysis, only the contigs that can be classified using a specific workflow and that are of interest to the scientific study are typically submitted to sequence repositories such as GenBank. The current approaches used for metagenomics extensively rely on similarity searches to known organisms and proteins, thus, suffe rs from the street light effect i.e. observational bias which occurs when people only search for something where it is easier to look. However, in a typical metagenomic data set, a range of assembled contigs cannot be functionally or taxonomically classified, a large proportion of which, even after excluding spurious contigs, bear no functional or sequence similarity to any known sequences and are often referred to as unknown or ‘dark’ sequence matter (Youle et al., 2012; Rinke et al., 2013; Krishnamurthy and Wang, 2017; Bernard et al., 2018). Although the terminology itself has been controversial (Bernard et al., 2018; Murat A., 2020), it typically refers to the sequences of unidentified taxonomic and/or functional origin (figure 1(b)). Generally, these unknown contigs (UCs) are excluded from downstream analyses. However, a number of recent studies have highlighted the importance of identification and categorisation of such unknown sequences.

Characterisation of metagenomically assembled genomes (MAGs) as microbial origin has strengthened the hypothesis that uncharacterised biological sequence matter is highly likely to belong to uncultured or unculturable bacteria, archaea and viruses present in the microbiome sampled (Rinke et al., 2013; Bernard et al., 2018; Thomas and Segata, 2019; Woyke et al., 2019). A study by Almeida, Alex L. Mitchell, et al. (2019) mined over 11,850 human gut microbiome data sets and identified nearly 2000 novel uncultured bacterial species from 92,143 genomes assembled from metagenomics data sets. Similarly, another focusing on multiple human biomes assembled 150,000 microbial genomes from 9428 metagenomic data sets (Pasolli, Asnicar, et al., 2019). The MAGs generated from these studies were consolidated to create a unified catalogue of 204,938 gut microbiome reference genomes (Almeida, Nayfach, et al., 2020). A range of different data mining studies have led to identification of novel microbes, including identification of novel bacterial and archaeal phyla and superphyla (Rinke et al., 2013; Saw et al., 2015).

Previous studies have shown that sequences of unknown lineage and unknown functions tend to be of viral origin (Youle et al., 2012). For example, a computationally identified phage crAssphage has been shown to constitute approximately 1.7% of all fecal metagenomic sequences (Dutilh et al., 2014). A study by Roux, Hallam, et al. (2015) mined 14,977 publicly available bacterial and archaeal genomes and identified 12,498 completely novel viral genomes linked to their hosts. Kowarsky et al. (2017) found that 1% of cell-free DNA sequences appear to be of non-human origin in human blood samples and only a small fraction of them can be mapped to currently known microbial sequences. Despite this, characterisation of unknown sequences in publicly available data repositories remains an on-going challenge in microbiome research (Thomas and Segata, 2019) and the identification of viruses in such UCs remain an even greater challenge due to the absence of a universal gene signatures and the high diversity in virus genome content (Wang, 2020). Overall, this highlights the widespread existence of potentially novel viruses and bacteria in the currently available sequence data sets, and that a systematic method to identify and catalogue them, especially in human data sets, would be extremely useful. The European Bioinformatic Institute (EBI) has developed MGnify (Alex L Mitchell, Scheremetjew, et al., 2018; Alex L Mitchell, Almeida, et al., 2019) that enables researcher to analyse their data using a standard metagenomic workflow. Similarly, there have been other community initiatives developed to forward this field of research (Pasolli, Schiffer, et al., 2017; Sczyrba et al., 2017; Alex L Mitchell, Scheremetjew, et al., 2018; Von Meijenfeldt et al., 2019; Paez-Espino et al., 2019; Tisza et al., 2020; Galloway-Peña and Hanson, 2020) however, a comprehensive computational framework and associated database aimed at identifying and quantifying UCs in different microbiome samples is still to be designed.

In this study, 1) we develop a framework to quantify the unknown sequence matter in human metagenomic data sets; 2) we compare the unknown sequences between samples, studies and microbiomes to determine whether these sequences are likely to be of biological origin and whether they are broadly distributed and 3) we compare the unknown contigs to currently known sequences in GenBank over the period of the study to determine the rate at which these unknown contig sequences are being taxonomically classified. All unknown sequences and associated metadata have been made publicly available for the research community and the original submitter.

## Methods

This study includes the data sets available within the EBI MGnify resource. All human microbiome studies submitted to ENA which were included in the MGnify databases were downloaded with the corresponding metadata on 19 April 2019. In order to obtain detailed metadata, each study was linked to the corresponding SRA repository using NCBI e-utilities (Sayers, 2018). As the focus was on shotgun metagenomic data sets, studies targeting metabarcoding-based sequencing methods such as 16S and amplicon sequencing were excluded as well as studies that solely focused on third party annotation i.e. analysis of previously published data and lack primary data were also excluded. In order to reduce sequencing technology related bias, the studies that utilised sequencing platforms other than Illumina were excluded. Very large studies involving >100 samples were discounted in order to get a cross section of different human microbiomes and geographical locations whilst keeping the overall data set size manageable. The filtered set initially comprised 44 distinct studies with 1130 samples of which 1121 were available to download. A script that uses parallel-fastq-dump (Valieris R., 2020) was developed to download reads in fastq format. In total, 1121 samples (789 paired-end [PE], 332 single-end [SE]) from 43 distinct studies were successfully downloaded and submitted to the pipeline. Out of 1121 samples, 158 could not be assembled due to insufficient reads and were excluded from downstream analysis (see supplementary method). In summary, 963 (784 PE, 179 SE) samples from 40 distinct studies were included and were processed using the complete metagenomic analyses pipeline described below.

This study set included a range of different sample types as described in the figure 3(a). It is important to note that this set is highly skewed towards the human gut metagenome that is normally sampled through fecal material and the oral microbiome was the second most common sample type included in the study. Although other metagenomes were under-represented, our study covered a wide range of samples from various human bodily sites and fluids. A miscellaneous metagenome labelled only as ‘Human’ was included in this data set that represents 3 distinct studies including PRJEB14301 (CSF, n=1), PRJEB21827 (A/B testing for colon model, n=12) and PRJEB6045 (metagenomics of medieval human remains from Sardinia, n=1).

**Figure 3:**
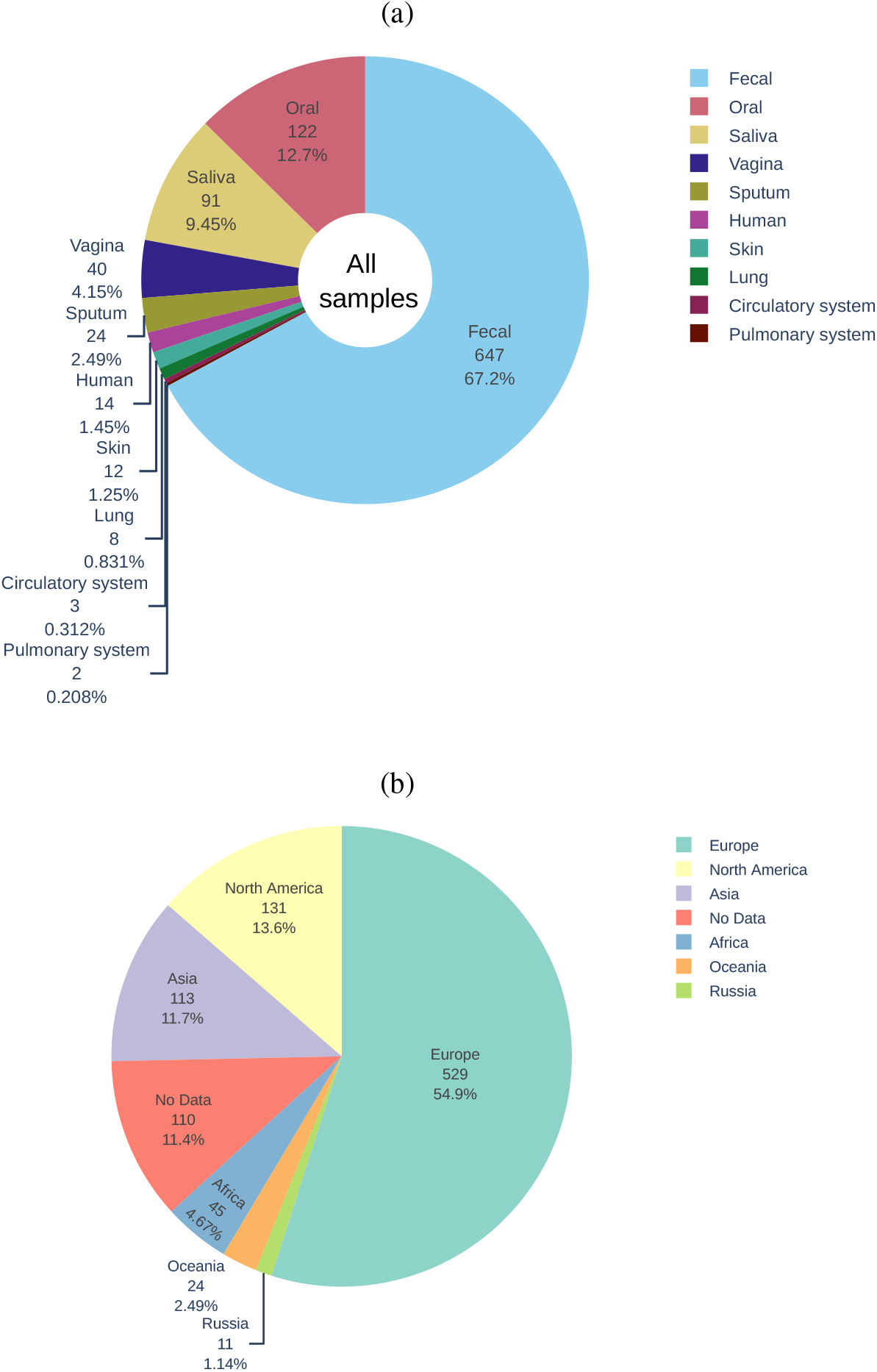
An overview of the human microbiome data set included in this study. (a) Distribution of samples included in this study per biome (n=963). (b) Overview of the geographical distribution of the samples included in the study (n=861) coloured according to distinct microbiome. The size of the slice represents the number and the proportion of samples.

In order to assess the quality of the samples and remove sequencing adaptors, all samples were processed through BBDuk from BBTools package (Bushnell B., 2019). BBDuk auto-detected the presence of the relevant adapter sequences from the input files specified and trimmed them. Additionally, commonly known sequencing contamination and spike-in sequences were also removed as part of this QC step. All reads that pass QC were retained and mapped to the human genome sequence build GRCh38 using the Burrows-Wheeler Aligner (BWA) (H. Li and Durbin, 2009), and unmapped reads were subsequently extracted using SAMTools (Heng Li et al., 2009). BBNorm (Bushnell B., 2019) was used to normalise reads based on the kmer coverage composition with a kmer threshold of 3 (`mindepth=3`). This step also enabled the acceleration of the assembly process as only a subset of reads were used to build the *de novo* assembly and resulted in better assembly quality overall (Crusoe et al., 2015). The read lengths varied widely between the samples and the studies, thus it was not possible to compare the quality metrics using the read-based measures as it would be misleading. To enable a comparison, quality assessment metrics were carried out for number of bases.

### *De novo* assembly and taxonomy label assignment

The normalised reads were *de novo* assembled using the SPAdes (Nurk et al., 2013) assembly pipeline, with the default parameters. A script was developed to extract contigs that were longer than 300 bases as short contigs do not contain a lot of information and they were excluded from downstream analysis as a precautionary measure. Although the normalised subset of reads was used to generate assemblies, these reads cannot be used to assess the assembly quality as they represent a small subset of the actual reads. To assess the assembly quality, the complete set of reads that did not map to the human genome were mapped onto the *de novo* assembled contigs with BWA (H. Li and Durbin, 2009) using the default parameters. The assembly quality statistics such as coverage, length, number of mapped reads were generated for each contig using `pileup.sh` from BBTools package (Bushnell B., 2019).

Contigs were searched against the GenBank non-redundant (nr) protein databases using the BLASTX algorithm implemented in DIAMOND (Buchfink et al., 2014). It carries out a six frame translation of the nucleotide sequences and then searches those translated sequences against the nr protein databases. This step enables the identification of distantly related homologues of the currently known sequences. The top 25 hits for each contig were extracted and analysed downstream (`--evalue=0.001`). The contigs that did not have any protein match were extracted and searched against the GenBank comprehensive nucleotide database (nt) using BLASTN (`--evalue=0.001`). This step helped to identify and remove non-coding sequences such as ribosomal RNA and untranslated regions of currently sequenced organism included in the databases.

To identify the geographical distribution of the raw data, location data was mined from the SRA metadata resources using pysradb (Choudhary, 2019) for each study. Geo-location information was available for 861 samples as shown in figure 4. A complete list of study location is shown in the appendix table S4. These samples were sequenced in various sequencing facilities across the world, and a complete distribution of the sequencing centre is shown in appendix figure S7.

**Figure 4:**
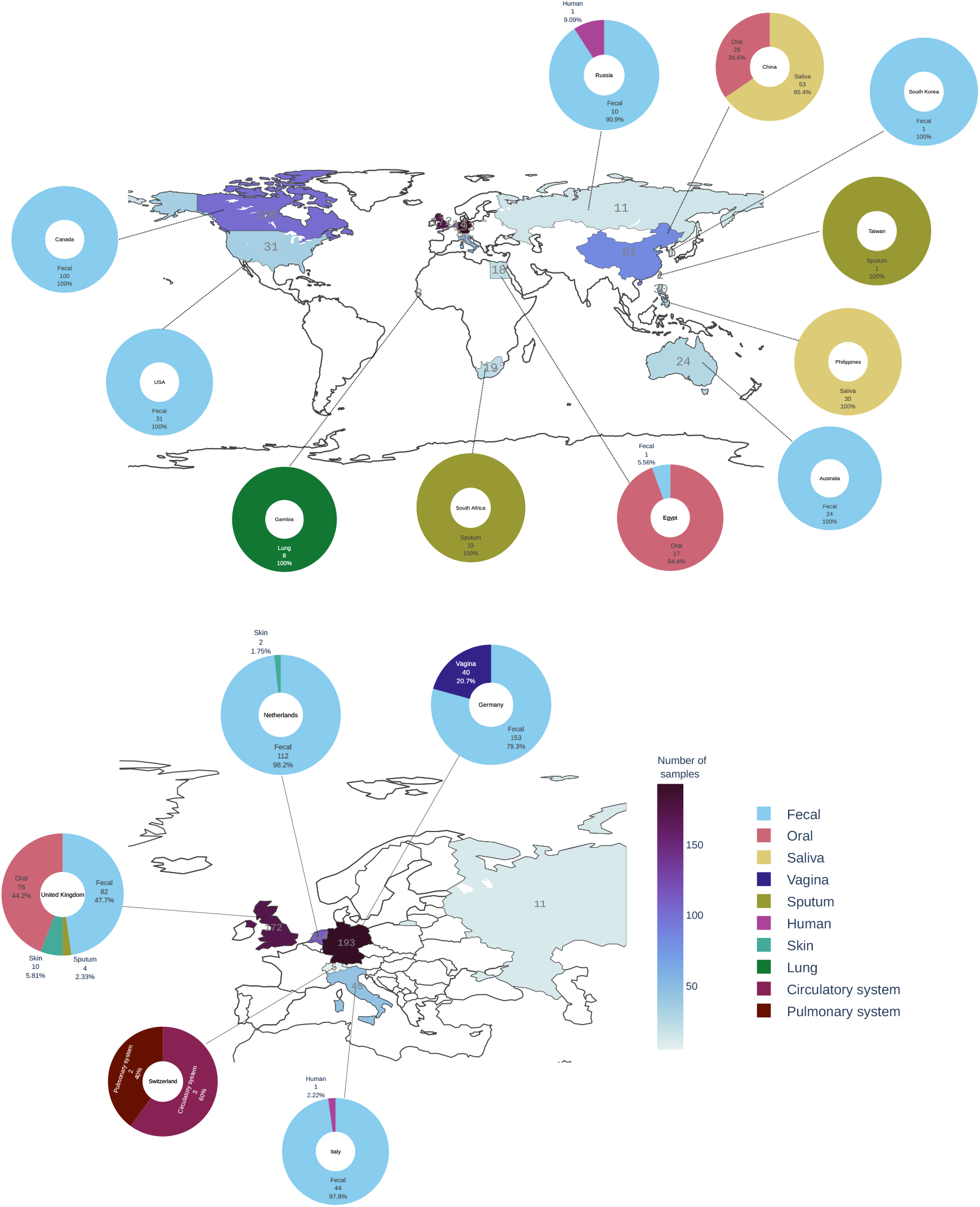
The geographical distribution of the human microbiome samples included in this study (n=861). Geographic locations are coloured according to the number of samples with darker shades representing the higher number of samples analysed. Samples originated from each location are represented by doughnut chart. Each doughnut is coloured according to microbiome and its proportion is represented by the slice of the doughnut.

### Unassembled sequences

‘Unassembled’ bases are defined as bases from reads that did not map to the human genome and could not be assembled into contigs. These were calculated from reads that did not map to assembled contigs. These bases/reads could not be classified as part of this project but were quantified as shown in the figure S8(b) - grey bars. Our quantification suggests that almost all microbiome samples have a proportion of unassembled sequences and on the sample the average values for this is around 23.91% (std: 26.59%). This unassembled sequences proportion was very high for samples originated from PRJEB15334 (mean: 51.17%, max: 97.67%, std: 24.30%) and PRJEB17784 (mean: 82.59%, max: 98.83%, std: 17.84%). Overall, 8.18% of all data fell into this category as described in figure S8(a). A range of possibilities from degraded nucleic acid to sequencing protocols could lead to poor quality data that cannot be used for *de novo* assembly.

### Control samples

The Human Microbiome Project mock community samples(n=9) were downloaded for study PRJNA298489 and were analysed using the metagenomic framework described above for quality control and workflow assessment. This would also allows us to validate the metagenomic analyses pipeline for this study.

### Post metagenomic analysis

All unknown contigs (UCs) were analysed further to get insights into the coding potential of those sequences. `getorf` tool from EMBOSS (Rice et al., 2000) suite was used to generate open reading frames (ORFs) from contigs (`-find 1, -minsize 300`) using the standard genetic code. These ORFs were searched against a range of different domains and functional identification databases included in the InterProScan.

To explore the sequence similarity between samples and the diversity of the unknown sequences, a nucleotide-based sequence similarity clustering which also used coverage was carried out using MMSeqs2 (Steinegger and Söding, 2017; Steinegger and Söding, 2018). All sequences with at least 90% sequence identity and at least 80% overlap were clustered using the MMSeqs2 `easy-cluster` pipeline (Steinegger and Söding, 2018). All UCs were processed through CheckV (Nayfach et al., 2020) pipeline to identify the UCs that were likely to belong to viruses. The most widely applied sequence similarity-based approaches rely on static versions of the databases to carry out the classification step of the analysis. In this study, the sequence databases utilised were downloaded on the 18th of April 2019. All results included in the study are based on the searches against this static version of the databases. However, the sequence database is ever expanding with new sequences being added to the databases each day. With newer sequences being added to these databases, it is very likely that unknown sequences transitions into the “known sequence space” over time. In order to identify the proportion of the unknown sequences classified over the period of the study, 4 distinct time points were considered. Static versions of the databases were downloaded on 31 October 2019, 5 March 2020 and 14 October 2020.

## Results

To quantify the presence of unknown sequences in human metagenomes, data sets included in the EBI MGnify (Alex L Mitchell, Scheremetjew, et al., 2018; Alex L Mitchell, Almeida, et al., 2019) were filtered to select for metagenomic data sets sequenced on Illumina platform (see Methods). A set of 963 samples from forty studies covering ten different microbiomes were downloaded from SRA repositories and analysed using the framework described in the Methods in order to characterise and quantify the unknown sequences in these samples. The studies included a total of 2.08 × 10^12^ bases of raw sequence data that was derived from a range of human microbiome studies including the following microbiomes (figure 3(a)): (1) circulatory system (n=2) (2) fecal (n=20) (3) lung (n=1) (4) oral (n=5) (5) pulmonary system (n=1) (6) saliva (n=3) (7) skin (n=2) (8) sputum (n=2) (9) vagina (n=1) and (10) human (n=3; miscellaneous). Geo-location information available for 861 of these samples shows that the data sets are globally distributed, but skewed towards western Europe (figure 4 and figure 3(b)). All samples were individually processed through the metagenomic analysis framework designed in this study (see Methods). The framework included an individual sample-based *de novo* assembly step resulting in a total of 44,238,374 *de novo* assembled contigs, 28,505,777 of them were longer than 300 nucleotide. Out of this set, 7,155,624 contigs were at least 1 kb long, 970,507 were at least 5 kb and 415,719 were at least 10kb long. The largest assembled contig was 1,380,230 bases long and was found in the human gut microbiome sample ERR505090. These contigs were then systematically processed by our metagenomic framework for BLASTX sequence similarity classification against the GenBank non-redundant protein database. Sequence similarity thresholds were used to sort the contigs into three classes: known (>80% similarity to a known protein sequence), partially known (>0 and <80% similarity to a known protein sequence), and unknown (no similarity to any existing sequence).

In total, 25,148,829 (88.22%) contigs were classified as known contigs whilst 2,517,700 (8.83%) of all analysed contigs were classified as partially known. The remaining sequences, referred to as unknown contigs (UC), are sequences that did not bear significant similarity to known sequence in the databases. Overall, 651,529 (2.29%) of contigs did not match any currently known sequences using our approach and were categorised as UCs. On average 1.3% of assembled bases per sample were found to be unknown. The proportion of unknown varied significantly between different assembled metagenomes as shown in the figure 5(a). Samples from some microbiomes such as the circulatory system did not contain any unknown sequences compared to the skin microbiome where this proportion was up to 25.85% for some samples.

**Figure 5:**
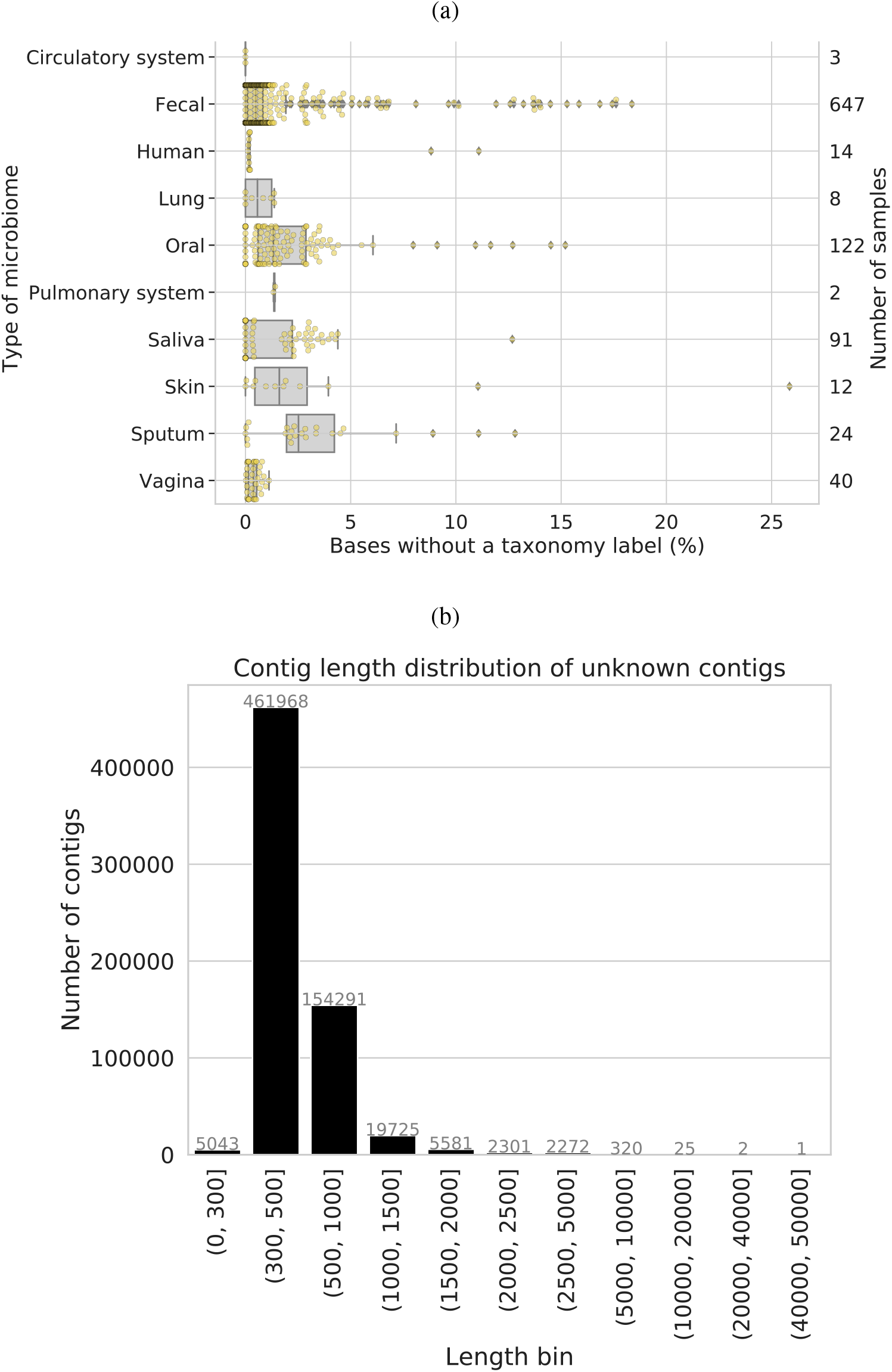
Quantification of unknown sequences in different human microbiomes. (a) Proportion of unknown bases in different human microbiomes. The proportion of unknown bases calculated from the unknown contigs for each microbiome. The secondary Y-axis shows the number of samples analysed in each category. Each individual sample is overlayed on the boxplot and are represented by small yellow circles. (b) The distribution of all unknown contigs in ten distinct length category. Each bar represent the contig length range shown on the X-axis with the number of contigs in the given category shown on the Y-axis and the exact number of contigs shown at the top of the bar.

The UCs varied largely in length and most of the UCs were 300-1000 nucleotides long (figure 5(b)). 95.36% (n=621,302) of all UCs were shorter than 1 kb and 4.59% (n=29,879) UCs were between 1-5 kb long. A set of 320 UCs fell within the 5-10 kb length category and 28 UCs were >10kb long. The largest UCs was 42.3 kb long and the second largest UCs was 21.3 kb long. A complete distribution of UCs across different miocrobiomes is shown in the figure S1 that shows that the largest UCs were assembled from fecal, oral and saliva microbiome.

To understand the coding potential of the unknown sequences, open reading frames (ORFs) were predicted. 273,590 ORFs that were at least 100 amino acid in length were generated using the standard genetic code. These ORFs originated from 215,985 distinct UC, showing that 33.15% of all UCs contained large ORFs. On average, ORFs were 157 amino acid (AA) long with a standard deviation of 87 AA residues. The longest ORF was 6,898 AA long. This set also included 2,713 ORFs with length of at least 500 AA and 256 that were at least 1000 AA long.

A detailed protein domain analysis for these ORFs was carried out using the InterProScan (Alex L Mitchell, Attwood, et al., 2018) protein analysis software. This tool searches the domain and functional signature of amino acid sequences against a range of distinct domain databases including Pfam (El-Gebali et al., 2018), CDD (Lu et al., 2019) and SUPERFAMILY (Gough et al., 2001). 36,354 ORFs orignating from 35,760 UCs could be functionally annotated using the InterProScan analyses, this number excludes hits to MobiDBLite and Coils databases as they predict disordered regions and coils structure of predicted ORFs as opposed to the domain signatures. An overview of the number of hits found to various InterProScan databases for each biome is shown in the figure S2. The most number of hits were found in the MobiDBlite (Necci et al., 2017) - a database that can predict the intrinsic disorder regions in the proteins. Overall, 5.49% of UCs (n=35,760) contained ORFs (n=36,354) with at least one identifiable domain. The functional classification of the ORFs was prominently centred around the Pfam database resource (El-Gebali et al., 2018). Pfam databases facilitate the domain-based searches against the set of protein sequences using Hidden Markov Model profiles. These types of searches can identify distantly related protein sequences. 16,839 ORFs originating from 16,705 UCs were found to match at least one Pfam entry and in total, 27,025 Pfam hits were derived (figure S2). All Pfam entries were collapsed down to their corresponding protein clans (grouping of related protein families) by mapping the Pfam IDs back to their clan membership. Figure S3 shows a heatmap of top 50 Pfam clans with hits to UCs ORFs predicted in different metagenomes. The most abundant hits were identified to clans tetratrico peptide repeat superfamily and leucin rich repeats. The largest number of hits was found in the fecal microbiome due to the high number of fecal microbiomes included in this study. Additionally, a range of other protein clans including those that represent Helix-turn-helix, beta-strands, polymerase and nuclease proteins were also found in this set. These results illustrate that the UCs sequences have known protein domains suggesting that these unknown sequences are functional and belong to organisms that are not yet fully sequenced or taxonomically classified.

### Unknown sequence clustering

To investigate the extent of sequence diversity and to identify UCs sequences present in multiple samples and microbiomes, sequence clustering was performed. MMSeqs2 (Steinegger and Söding, 2018) generated 464,181 clusters of which 377,855 were singletons i.e. did not cluster with any other sequences. These singletons were excluded from the cluster analysis described below. 86,326 clusters comprised two or more sequences with mean cluster size of 5.7 contigs and a standard deviation of 8.1. The largest cluster contained 153 sequences which originated from the fecal microbiome from 8 distinct bioprojects (figure S6(c)). A cluster size distribution across different microbiomes is shown in figure 6 and a detailed cluster size distribution with cluster representative length is shown in the figure S4. 89.42% of 273,674 UCs (n=244,730) were clustered into single microbiome clusters, 10.58% UCs (n=28,944) were found in clusters that contained sequences from two or more microbiomes. To compare that with specific studies, 39.4% UCs were clustered into bioproject specific clusters and the remaining 60.6% UCs (n=165,851) were grouped into clusters originating from two of more bioprojects. 78,139 (90.52%) clusters contained sequences from a single microbiome and 7,645 (8.86%) clusters included sequences from two microbiomes. Only a few clusters were comprised of members from 3 (n=512) or 4 (n=30) microbiomes. The largest multi-microbiome cluster contained 57 sequences (304-9,080 bases long) from 4 distinct microbiomes and bioprojects and contigs assembled from 12 samples. The largest single microbiome cluster contained 153 sequences (6,640-300 bases long) from fecal microbiomes with contigs assembled from 46 distinct samples covering 8 different studies. Overall, this clustering method produced very small, study specific clusters. A set of 464,181 UCs was obtained by combining the cluster representative sequences with the unclustered singleton UCs and used to determine the rate at which UCs are classified.

**Figure 6:**
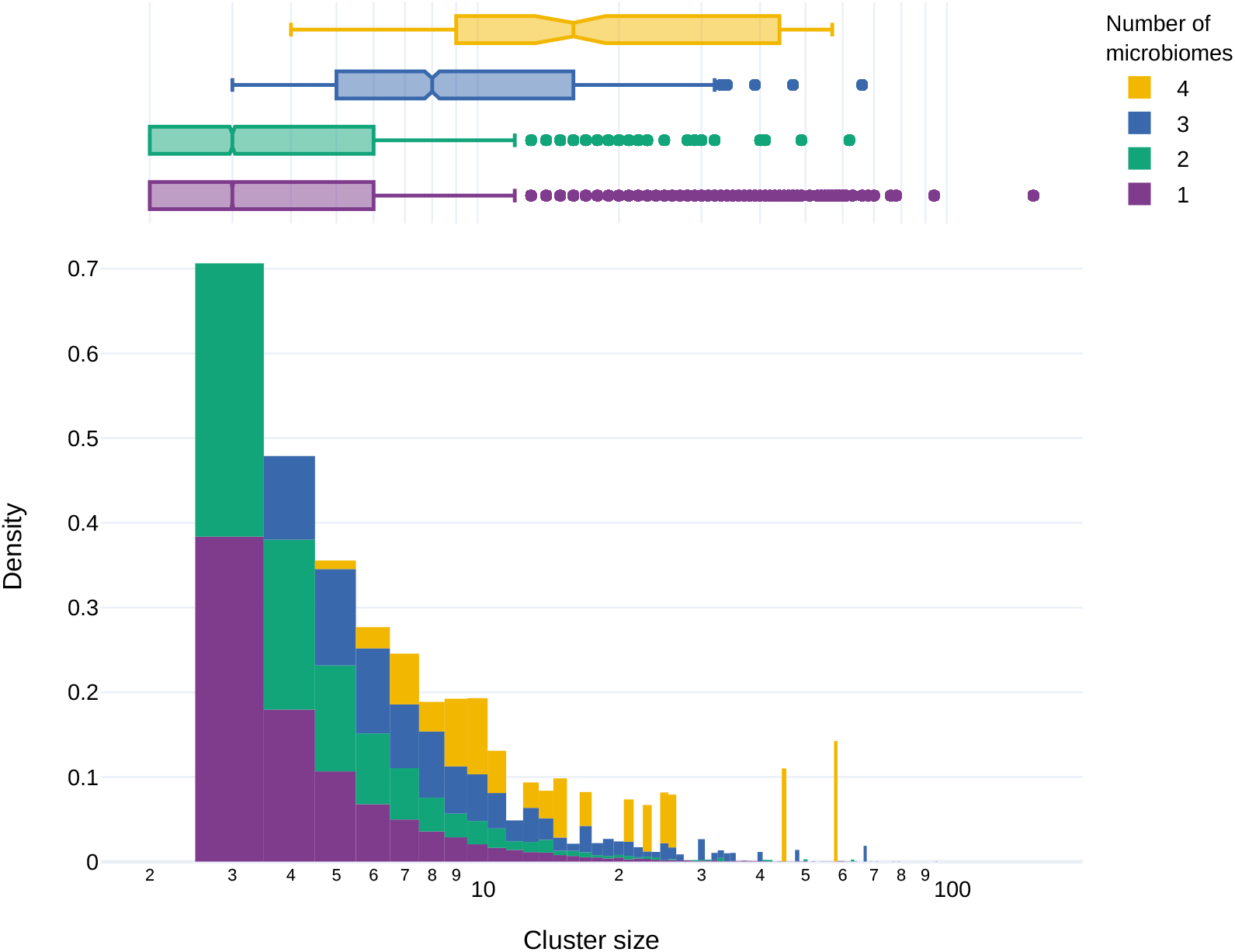
Distribution of cluster sizes and presence of cluster members across different microbiomes.

### Classification of unknown over time

In this framework, the unknown sequence identification is dependent on the publicly available nucleotide or protein sequence databases. These data repositories are updated regularly with new sequence data being deposited from around the world. However, typically, the sequence searches are carried out against static versions of the databases. Our analysis conducted against the databases downloaded on 18 April 2019 identified 651,529 UCs that were collapsed down to a set of 464,181 UCs following the cluster analysis. Subsequent analyses on 31 October 2019 and 5 March 2020 produced a set of 613,726 and 558,711 UCs respectively. The final number of sequences that still lacked a taxonomy label was down to 459,147 after the most recent analysis carried out against the databases downloaded on 14 October 2020. 29.5% (n=192,382) of the sequences compared to the initial set of unknown matched to at least one sequence from the updated databases in the BLASTX and the BLASTN steps of the analysis. Similarly, 27.6% (n=128,288) of the representative set sequences could be labelled taxonomically with the updated databases. A rate of taxonomic characterisation of 1.64% of unknown sequences being characterised per month was calculated from the complete set. This rate was estimated to be 1.54% for the representative set. Moreover, as shown in the supplementary figure S5, a range of long UCs still remained unknown even after the similarity sequence-based analysis carried out on the 14 Oct 2020.

From a set of 192,382 contigs that were labelled taxonomically after the most recent analyses carried out on 14 Oct 2020, 167,864 were identified using BLASTX and 24,518 were identified using BLASTN. 106,739 UCs from the BLASTX classified set were categorised as known and 61,125 contigs were categorised as partially known. A large majority of these contigs (97.11%, n=162,987) were also deemed to be bacterial. The remaining contigs were divided between cellular organisms (n=2,104), archaea (n=930), viruses (n=858), root (n=827) and eurkaryota (n=140). 76.55% of all BLASTN hits were matching to bacteria (n=18,768), 17.88% matched to viruses (n=4,383), 1.99% matched to eukaryota (n=487) and 0.03% archaea (n=7). The hits that could not be mapped to a superkingdom and were divided between unidentified plasmid (n=544), root (n=294), cellular organisms (n=20), uncultured organism (n=14) and synthetic construct (n=1). These results reiterate our initial hypothesis that the majority of UCs represent currently unknown microbial genomes.

### Predicted viral contigs

195 UCs were shown to contain a virus specific functional domain which was parsed using the term ‘virus’ or ‘viral’ in the interproscan analysis signature description column. Results with the term ‘phage’ were not included in this subset as a range of phage domains are also present in the host bacterial genomes. These domains were predominantly identified using the Pfam (n=125) analysis. The most abundant virus specific domain was Vaccinia Virus protein VP39 and it was found in 53 UCs derived from fecal (n=23), saliva (n=14), oral (n=12), sputum (n=1) and human (n=3) microbiomes and it was identified by Gene3D analysis. The largest UCs containing this domain was 3,661 bases long and was found in sample ERR1474567. Another frequently found domain in the UCs was podovirus DNA encapsidation protein Gp16 domain. It was found in 25 UC, out of this set 23 UCs were assembled from fecal microbiome. The largest UCs containing this virus specific domain was 9 kb long contig shown in figure 8(a) assembled from PRJEB18265. This UCs was clustered with 24 other sequences (See Unknown sequence clustering) that were assembled from 11 samples representing 5 distinct fecal microbiome studies. These results indicates that this UCs represents a completely novel genome of virus that is likely related to currently known podoviruses.

**Figure 7:**
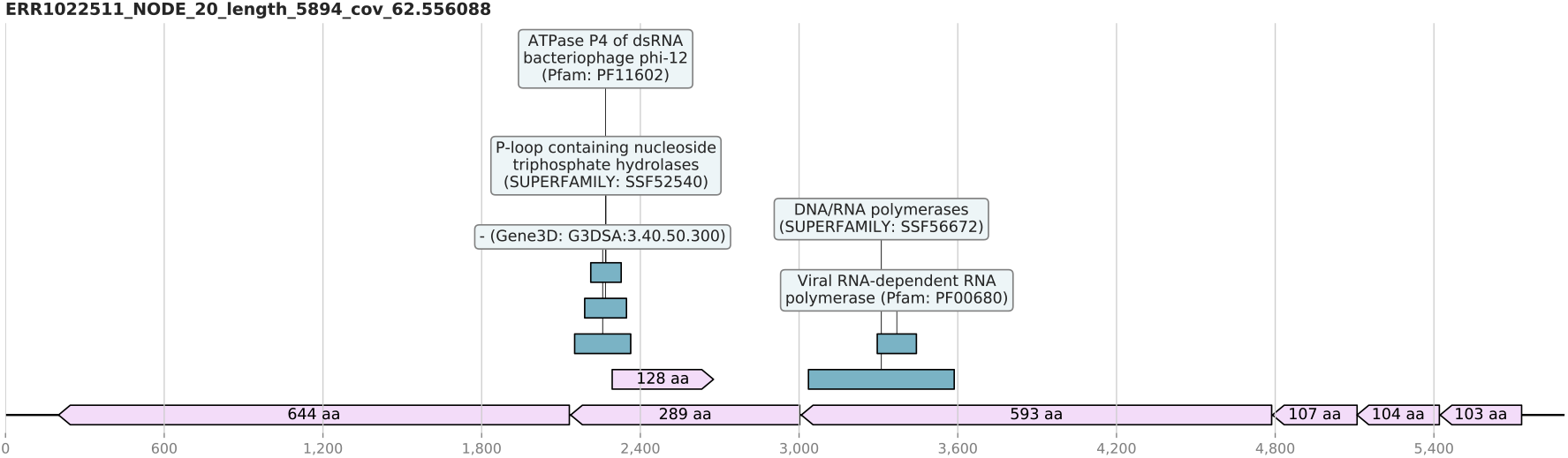
A potential novel dsRNA phage segment found among the UC set that is speculated to be related to currently known Cystoviruses.

**Figure 8:**
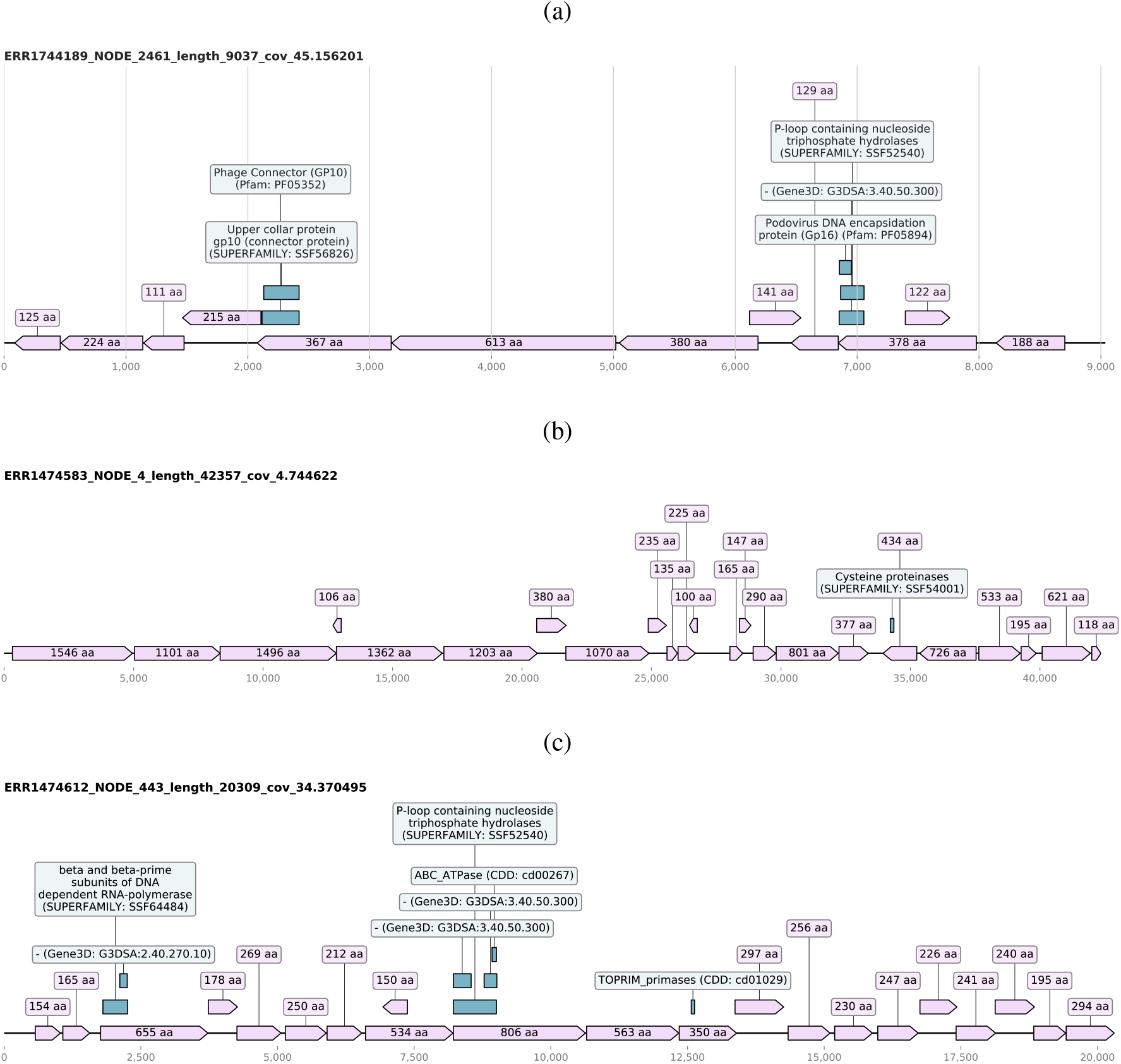
The genome diagrams of large unknown contigs showing the open reading frames (ORFs) in light pink shade with the ORFs lengths as their corresponding labels and the green boxes illustrating the InterProScan computed presence of domain signature. (a) The largest contig with podovirus DNA encapsidation protein Gp16 domain. (b) The largest unknown contig assembled in the set that is categorised as unknown even after the most recent similarity-based search at in 14 Oct 2020. (c) Unknown contig of length 20,309 bases that was described to contain a range of domains including a potentially virus specific RNA polymerase domain.

The largest UCs containing a viral RNA dependent RNA polymerase (Pfam:PF00680) domain was found in the sputum microbiome sample ERR1022511. This UCs was 5,894 bases long and contained seven ORFs that were at least 100 AA long (figure 7). A 269 AA long ORF contained ATPase P4 of dsRNA bacteriophage phi-12 (Pfam:PF11602) domain suggesting that this UCs represents the large segment of a novel double-stranded RNA phage which are usually categorised in the virus family *Cystoviridae*. The genomes of these phages are composed of three linear dsRNA segments with a total genome length of 12.7–15.0 kbp and all segments code for various proteins (Poranen and Mäntynen, 2017). Although several other UCs were found in the same sample, none of them displayed any sequence or functional similarity to the other two segments i.e. small and medium segments of Cystoviruses. However, UCs that could potentially belong to novel cystovirus-like genomes were extracted based on the sequence length, GC content and sequencing depth criteria. Moreover, this UCs representing a potentially novel relative cystoviruses did not match to any known protein or nucleotide sequences even in the most recent analyses confirming the discovery of a novel virus.

### The large unknown contigs

The largest UCs was assembled from the saliva sample ERR1474583 and was 42,357 bases long. This contig did not cluster with any other contigs and has 23 ORFs that were over 100 AA long. One of the ORFs that is 434 AA long comprised of the cysteine proteinases domain (SUPERFAMILY: SSF54001) according to the InterProScan analysis. This contig still remained unknown after searches against the most recent version of the databases suggesting that the organism this genomic sequence belongs to is still to be identified and fully sequenced. A snapshot of the ORFs and domain is shown in figure 8(b), highlighting the presence of coding regions across the entire length of the UCs sequence. Based on the results we have obtained here, we predict that this UCs sequence is likely to be of microbial origin as it lacks non-coding region. CheckV analysis predicted it to be a viral genome fragment with the presence of two identifiable viral genes albeit with low quality as per the MIUVIG (Roux, Adriaenssens, et al., 2019) standards due to the lack of similarity to any known sequences. This strongly suggests that this UCs can potentially be a representative or partial genome sequence of currently unknown and completely novel virus.

A 20,309 nucleotide long contig from saliva sample ERR1474612 clustered with two very short contigs from the same study. As shown in figure 8(c), long ORFs were predicted across the whole sequence. Some of the predicted ORFs were found to have interesting domain signatures (figure 8(c)). An ORF that is 655 AA long shows the presence of DNA dependent RNA polymerase domain (SUPERFAMILY: SSF64484). A CheckV (Nayfach et al., 2020) analysis of the contig also predicted it to be of viral genomic origin, however, it was predicted to be an incomplete genome. This UCs was shown to have very low identity (<30% sequence identity with 2% of query coverage) to hypothetical protein of Firmicutes bacterium (HAB66316.1) and AAA family ATPase from Sharpea azabuensis (23% sequence similarity). When the evalue threshold was removed, a total of 8 BLAST hits were obtained and 3 out of 8 hits were to a range of phages including Bacillus phage vB_BpuM-BpSp, Vibrio phage 2 TSL-2019 and Ralstonia phage RP12. These hits ranges from hypothetical and putative proteins. All these matches were localised to a short regions between 8,217-8,915 which was shown to contained ATPase and P-loop containing nucleotide triphosphate hydrolases domains (figure 8(c)). Notably, no nucleotide sequence hits were identified for this UCs. Although these results have bacterial hits, it is likely that this UCs represent a complete of partial genome of a novel phage that infects the host bacteria e.g. firmicutes.

### Short circular contigs

A range of circular contigs with direct terminal repeat (DTR) and inverted terminal repeat (ITR) signatures were identified using CheckV in the UCs data set. A total of 1,839 containing repeat signatures were predicted of which 1,771 contained DTR signatures and 68 contained ITR signatures. 94 of these UCs were at least 1 kb long suggesting circular genomes and 48 of them contained a range of 55 bases long terminal repeats. A cluster of 8 sequences from 2 different microbiomes and studies were identified to contain similar sequences (71-100 percent similarity) assembled from different samples (table 1). Four cluster members were 2,110 bases long, one sequence was 1,983 nucleotide long and the cluster representative was 3,165 nucleotide long. The cluster representative sequence contained multiple copies of the same ORFs suggesting the presence of multiple genome copies, sequencing error or miss-assembly. Most of these sequences contained a 50 bp long DTR sequence signature ‘GTGCATTTTTTTTGTGCACTTTTTCAAAAAAAC-CGTGAAAAAAATTCATT’. These contigs contained two distinct ORFs, which were 125 AA and 144 AA long. Similarly another 50 bases long DTR signature ‘AATGAATTTTTTTCACG-GTTTTTTTGAAAAAGTGCACAAAAAAAATGCAC’ was observed in another cluster that had 7 member sequences ranging in similarity from 31 to 100 percent and assembled from 7 distinct samples. All but one members were 1,770-1,771 bases long. These contigs also contained two ORFs that were 102 AA and 106 AA long. These ORFs did not match any existing protein sequences in the databases. These circular contigs were assembled from a range of oral microbiome samples from study PRJNA230363. Similarly, a range of contigs (n=9) that contained Inverted Terminal Repeats (ITR) were also identified in this data set. A cluster of 5 distinct circular contigs assembled from distinct samples from the fecal microbiome (PRJEB7949). Four out of five of these circular contig contained the ITR sequence ‘CGAAACGATTGCCCAGAGAGATGACTGTCAATCCGCCCGATTATTGGGCGCTTAC’. They also contained a 138 AA long ORF. These short circular UCs did not bear any sequence or functional similarity to known sequences or domains so their biological origin is difficult to predict. However, based on their genome organisation and size distribution, we predict that they are likely to represent either novel CRESS DNA virus groups or novel satellite virus-like groups.

**Table 1:** Circular contig clusters with direct and inverted terminal repeats

| Study ID(s) | Cluster size | Typical contig length | Repeat type | Sample type | Sequence similarity (min-max) |
| --- | --- | --- | --- | --- | --- |
| PRJEB14383;<br>PRJNA230363 | 8 | 2110 | DTR | Saliva; Oral | 71-100 |
| PRJNA230363 | 7 | 1771 | DTR | Oral | 31-100 |
| PRJEB7949 | 5 | 1337 | ITR | Fecal | 67-100 |

### Control samples

The HMP mock community samples (n=9) were downloaded for study PRJNA298489 and were analysed for quality control and workflow assessment. These are control samples which are not expected to yield UCs, but if they do, those UCs could be due to sequencing/assembly error or common lab contaminants. Out of the complete set, four HMP samples did not contain any UCs as expected whereas SRR2726666, SRR2726669 and SRR2726672 contained one UC each but their lengths were short varying from 323 to 449 bases. The remaining two samples SRR2726670 and SRR2726671 contained 28 and 18 UCs each. The largest UC assembled in the mock sample was 3,965 bases long and was found in SRR2726670, only 3 UCs were >= 1 kb. These UCs were searched against the most recent version of the databases downloaded on 14 Oct 2020 and only 8 short contigs; 4 from SRR2726670, 2 from SRR2726671 and one each from SRR2726666 and SRR2726669 remained in the UCs category. These remaining UCs were only 330-513 bases long. These results validate the UC analysis framework developed here and highlights that the even in control samples, there are a very minor number of short UCs to be found.

### Resources

All assembled unknown sequences generated here are submitted to ENA as third party annotations and are accessible under bioproject PRJEB41812. This allows the UCs data to be properly linked to its original samples and studies. Results and code generated in this study is available on https://github.com/sejmodha/UnXplore. A consistent data labelling scheme is utilised across all studies and sample. For traceability, all UCs fasta identifiers start with SRA sample identifier. All ORFs contain the exact same naming scheme with a suffix ‘_’ and ORF number starting with 1. A complete metadata table is provided to link any new sequence data to its corresponding bioproject and sample. Functional domain predictions and clustering results are annotated with relevant metadata and provided in a tabular format.

## Discussion

In this study, we have developed an automated framework that can systematically quantify the proportion of unknown contigs (UCs) in meta-omics samples. Whilst the presence of UCs is well recognised, this is the first study that addresses the question of UCs comprehensively and quantifies it across different human microbiomes. Our approach utilises sequence similarity-based taxonomic categorisation to identify the sequences that cannot be categorised. We define these UCs as the sequences that do not match known sequences in the databases with a predefined sequence similarity threshold of evalue 0.001 which is a very lenient threshold, anything with evalue higher than this is unlikely to truly be related to the database sequence hit. We show that on average 2.29% of assembled contigs are categorised as unknown in different human microbiome studies. Moreover, a subset of those with unknown sequences could be translated and contained protein domains, thus we were able to find functional similarity to 5.49% of taxonomically unknown contigs. We have generated a comprehensive catalogue of 651,529 UCs that do not bear any sequence similarity to sequences present in the widely used GenBank protein and nucleotide databases. Although sequence similarity based approaches are dependent on the databases, the protein sequence-based approached implemented here is highly effective in fishing out distantly related homologues of known sequences available in the databases (Altschul et al., 1990) and thus provides better resolution for sequence classification compared to those solely based on the genomic signature-based binning (Chen et al., 2020). This study highlights the importance of avoiding the “street light” effect i.e. observational bias arising from classifying metagenomic sequences on the basis of related sequences that already exist in the databases. Here, we have aimed to eliminate such observational bias by performing a comprehensive data mining of the human microbiome data and cataloguing the UCs, their abundance in different human microbiomes and their overlap between different samples.

This study has enabled the identification of a range of genomic sequences that are hypothesised to belong to currently uncharacterised organisms that are often found in similar samples and/or microbiomes. A range of large UCs with and without known protein domains are presented here. However, the complete set includes a large number of UCs that still remain unknown and can be mined further to study their biological origin. A third of all UCs (n=215,985) contained large predicted open reading frames (at least 100 amino acid long) that were predicted using the standard genetic code. Using alternative genetic codes may expand this set further by revealing novel, potentially different open reading frames generates from the UCs. A small proportion of these open reading frames contained domain signatures confirming the presence of currently unidentified organisms. Moreover, a comprehensive clustering analysis has led to the identification of UCs that were present across different human microbiomes (as well as from different samples/studies investigating the same human microbiome) indicating that we have discovered potentially widespread and as yet unclassified novel biological organisms within the human microbiome. The multi-microbiome clustering approach applied here provides an interesting way to understand the diversity and the distribution of the UCs across different microbiomes and geographical sites. Although it is impossible to identify the true clusters present in the data due to the novelty of the UCs, the clustering approach helps to identify obvious patterns of sequences similarity between microbiomes and studies. This approach provides an additional dimension by capturing unknown sequences that are shared between different projects or human microbiomes.

Short contigs i.e. those less than 1-5 kb are often ignored in most data mining and exploration research typically in studies that employ a contig binning step as binning has been shown to be less sensitive for short contigs (Breitwieser et al., 2018; Chen et al., 2020; Mallawaarachchi et al., 2020). The clustering and timepoint analyses carried out on short UCs has shown that these short UCs are originating from biological entities and predominantly represent the novel microbial sequences that are currently uncatalogued. This has been demonstrated with the example of short circular sequences with terminal repeats. Short contigs that are typically excluded from large microbiome mining studies employing the metagenomic binning approach but were studied in detail here. These short UCs are found across multiple human microbiomes and samples, we speculate that these are of viral origin and could potentially represent novel CRESS DNA or satellite viruses, although the ORFs originating from these genomes do not bear any sequence of functional similarity to the typical rep and cap genes. Moreover, a number of large contigs were found to contain various functional ORFs and domains often originating from virus or phages indicating that a proportion of UCs are very likely to be novel viruses that infect currently uncharacterised microbes. In our approach, we have implemented a protein sequence similarity-based identification that enable the identification of distantly related sequence homologues (Altschul et al., 1990). This approach can potentially ‘classify’ contigs of viruses or phages as their corresponding host with very low sequence similarity. Indeed, viruses are well known to mimic their host genomic signatures by incorporating genomic sequences from their host into their genome. We anticipate that the virus diversity described in this manuscript is reasonably underestimated due to this specific characteristic of viruses and speculate that a range of assembled contigs classified as bacterial with very low sequence similarity across a short genomic coverage are likely to be of virus origin. This hypothesise will need to be tested further by mining the ‘known’ and ‘partially known’ contigs systematically. The HMP control sample analyses resulted in only a few UCs validating the UC identification approach implemented in our framework. The results generated from this study can be extended to identify the organisms that co-occur in different microbiomes, which in turn can help to inform the interactions between these organisms and how it affects their hosts - humans. Despite having sequenced human microbiomes extensively, our understanding about how these microbes interact with humans remains limited. These large scale explorations can help to understand the human holobionts and the interactions of macro- and microorganisms. Based on these results, we do not know whether the microbes identified in different studies are consistently associated with human or they are just passing association captured at the time of sampling, the latter would make it even harder to make comparisons between samples and microbiomes.

The UCs landscape changes over time as more sequences get characterised and added to the ever expanding sequence repositories. This was demonstrated by comparing the UCs to different GenBank databases over the course of 18 months. We have estimated that 1.64% of the UCs identified in this study are getting characterised each month. However, this number would be highly dependent on the types of data deposited in the International Nucleotide Sequence Database Collaboration (INSDC) resources. This study provides a strong foundation of preliminary estimation of this rate and UCs would need to be analysed at multiple future time-points to determine how the rate at which the UCs are being classified, changes over time. Additionally, the time-point analysis also provides strong evidence of the real biological entities being assembled and characterised in our study. Indeed, a proportion of the UCs were taxonomically classified during the period of the study. This delineation of the UCs demonstrates that the unknown matter that surrounds us largely belongs to currently uncultured, unidentified microbes that we interact with on a daily basis. The technological advances have accelerated the speed at which genomic sequences belonging to novel uncultured organisms are being deposited in INSDC databases. This sharp increase of metagenomically assembled microbial genomes has led to the scientific community driving the development of genomic data and metadata standards such as MIMAG (for bacteria and archaea) (Bowers et al., 2017) and MIUVIG (for viruses) (Roux, Adriaenssens, et al., 2019) for consistency and comparison purposes. The taxonomic classification landscape has also faced a tectonic shift whereby it is moving from the phenotype-based classification to more holistic sequence-centric phylogenetic classification, e.g. GTDB (bacteria and archaea) (Parks et al., 2018) and ICTV (viruses) (Simmonds et al., 2017). These changes enable the incorporation of the uncultured sequence diversity into the microbial taxonomy and will provide a more comprehensive understanding of the complex phylogenetic relationships and interactions between different microbes.

The metagenomics analysis framework developed here works as a proof of concept for overcoming the challenge of the quantification of unknown in already ‘analysed’ data sets. The pipeline developed here is flexible and can be applied to any microbiome. To get a cross section of different human microbiomes and geographical locations whilst keeping the overall data set size manageable large studies involving >100 samples were discounted. This framework can readily be applied to routine metagenomic exploration, which can help to gain further understanding of the landscape of sequences of unknown origins. However, the framework applied here is easily portable to metatranscriptomics data. In fact, a couple of the bioprojects (PRJEB10919 and PRJEB21446) analysed in this study were indeed from a metatranscriptomic study. It is important to note that unlike other studies that often focus on the cross assembly of different samples, each sample was assembled individually here. This is regarded as best practice when a cocktail of samples from unrelated studies are analysed in bulk. The co-assembly would often lead to fragmented assembly as the complexity of sequences originating from multiple samples would be much higher compared to a single sample. On the contrary, independent assembly is expected to capture better diversity across each sample with high quality genomes assembled from each sample (Olm et al., 2017). Typically the sequence similarity-based approach is less reliable for unrelated sequences as the similarity search tools heavily rely on the databases used in the analysis. Like most other pipelines, this framework classifies the sequences with respect to a static version of the reference sequence databases. The search results are as good as the data in the ever expanding repositories that are often too large to be hosted on a local computer. In order to improve this, an alignment-free approach could be explored. The development of a general purpose alignment-free prediction method that can categorise the sequences based on the genomic composition would be suitable for the downstream analysis of the UCs. The UCs classification is highly dependent on the methods employed to identify and quantify the unknown. Moving away from the sequence similarity based methods would help to categorise and classify the currently unknown sequences better. Machine learning based approaches might be deemed suitable in certain circumstances to overcome the similarity threshold based approaches. In case of completely novel sequences that bears no similarity to currently known sequences, significantly rigorous training sets and features would need to be identified and be built into the models in order to make accurate predictions as machine learning approaches are highly reliant on the training data the models have been developed with.

This study demonstrates that there is a large diversity of unknown sequences embedded within various human meta-omic samples available in public repositories. It is clear that the unknown sequence landscape observed in this study is likely to be the tip of the iceberg, and, as we scan more microbiomes and extend this to less-studied environments e.g. insect metagenomes, we are likely to gather a better understanding of the unknown sequence space. As more species and environments are sequenced more readily, the rate at which the unknown sequences become known would also change. Our results of novel viruses indicate that the unknown microbes are likely to have different properties to those currently associated with the taxonomically classified organisms. Our study also shows that at least some of these unknown microorganisms are omnipresent and possibly thriving in nature. To overcome this, more comprehensive resources including searchable databases such as those enabled using BIGSI (Bradley et al., 2019) and federated indexes (Martí-Carreras et al., 2020) could be created for the unknown sequence data and metadata. This would allow researchers to explore the human metagenomic sequence space in more holistic manner and in turn provide a better understanding of microbial diversity interacting with and within human hosts. It would enable researchers to search, link and explore the unknown sequences present in different microbiomes, studies and samples. Such resources could help in speeding up the pace at which unknown sequences can be ‘classified’ and make it easier for researcher to determine the functional and/or ecological importance of the organisms the sequence comes from. A concerted efforts could help to pin down human-microbial interactions in a broader context such as linking unknown microbes to human diseases and disorders of unknown aetiologies.

## Acknowledgements

All authors would like to thank Prof Andrew Davison, Hilary Fawcett, Dr Quan Gu, Maha Maabar, Dr Josh Singer and Dr Sreenu Vattipally for contributing to an internal hackathon organised in 2016 where the idea of mining metagenomic data sets for identification of ‘dark matter’ was developed.

