## Supplementary material for "Quantifying and cataloguing unknown sequences within human microbiomes" (from another version of this preprint: version 1, 2021-01-22)

[#]Indicates last author

January 22, 2021

### Supplementary Data

### Supplementary text

#### CheckV predicted viral contigs

All UCs were processed through CheckV pipeline (Nayfach et al., 2020) to identify the UCs that were likely to belong to viruses. A total of 47,702 UCs were predicted to be of viral origin and 7,696 of them were at least 1 kb long. A set of 11,121 of these UCs were predicted to have at least one viral gene and 1,712 UCs of this subset were at least 1 kb long. These 1,712 UCs were mapped against the results of the most recently carried out BLASTX analysis for validation. 529 of the predicted viral contigs matched to bacterial protein sequences with low sequence identity with a mean percent identity of 48.43. However, these results are based on short protein sequence hits on bacterial proteins indicating that these UCs are likely to be phage genomic signature that matched to bacterial protein in absence of a phage sequence in the database specifically as protein-based similarity search are able to identify distantly related homologues of query sequences. These results can help to stipulate that the actual diversity of virus sequences presence in the UCs set is largely underestimated. It is highly likely that a range of contigs that match distantly related protein sequences of bacterial origin, are in fact derived from unknown and uncultured novel viruses, such as phages, that infect bacteria.

#### The large unknown contigs

The largest multi metagenome UCs (figure S6(a)), was assembled from SRR2037089 from the oral metagenome and was 14,958 bases long. It was clustered with 33 other contig sequence assembled from 12 distinct samples from oral (n=8;PRJNA230363), sputum (n=3;PRJEB10919) and saliva (n=23;PRJEB14383) microbiomes. These three distinct studies contained samples from distinct geographic locations: PRJNA230363 from China, PRJEB14383 from Philippines and PRJEB10919 from South Africa suggesting that this unknown organism is broadly distributed in its association with humans. The second largest member of this cluster was 9,791 bases long and was assembled from a separate sample (SRR2037087) from the same study. This large contig was deemed to be identical to the cluster representative. The largest cluster member from saliva microbbiome was 1.5 kb long and was assembled from ERR1474566. On the contrary, the contigs assembled from the sputum microbiomes were significantly smaller with length ranging between 479-533 bases, indicating the fragmented assembly and the presence of partial sequences.

A large contig of length 21,357 was identified in oral microbiome shown in figure S6(b). This contig was assembled from run ERR1611386 and was clustered with 16 other sequences from bioprojects PRJEB12831 and PRJEB15334. Other members of the clusters originated from 5 distinct samples and were between 306-6,109 bases long. This contig contained the largest predicted ORF that was 6,898 residues long. 14 out of the 16 other contigs within the cluster contained partial sequences belonging to this ORF. This contig also did not have a taxonomic homologue identified in any of the most recent similarity sequence-based searches. Additionally, the largest ORF was predicted to contain P-loop containing nucleoside triphosphate hylrolases (SUPERFAMILY: SSF52540) signatures.

The largest cluster contained 153 sequences (figure S6(c)) had a cluster representative that was 6,642 bases long assembled from sample ERR1297807 from PRJEB12357. This cluster representative was predicted to contain 9 distinct ORFs. The cluster contained 35 other sequences that were at least 1 kb long. Additionally, other contigs (6,015 bases long from ERR537012 and 5,344 bases long from ERR537011) from as separate study (PRJEB6542) were found in this large cluster.

### Supplementary figures

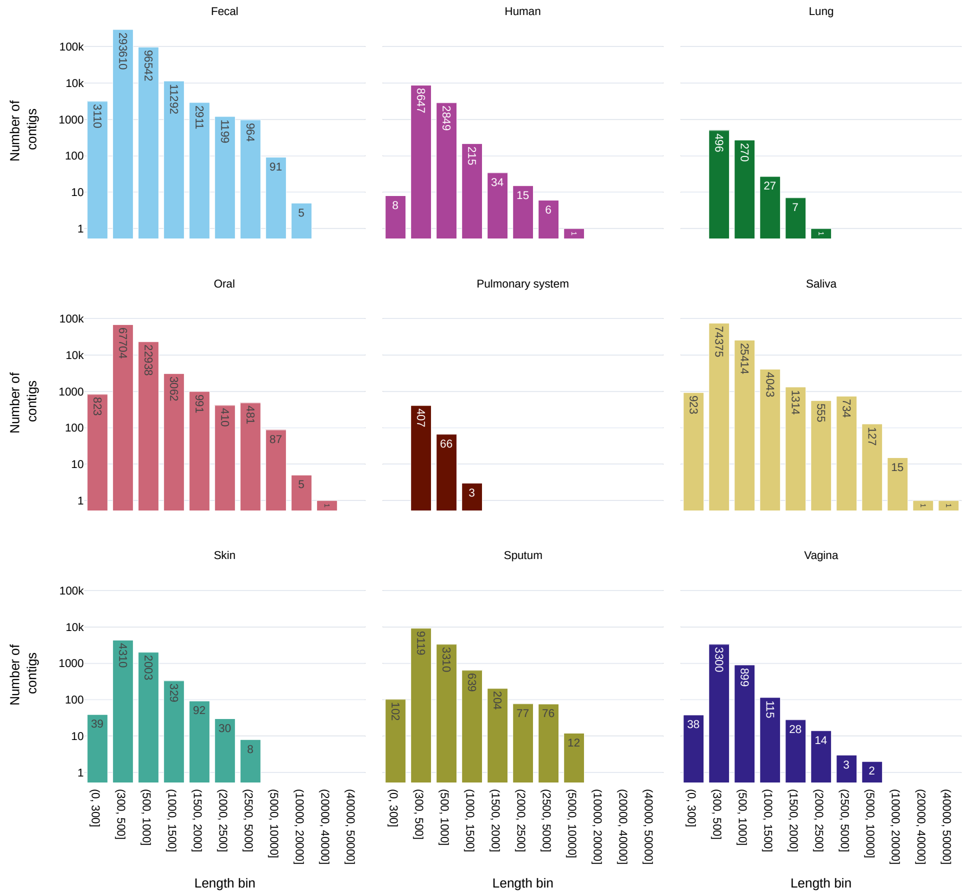


Figure S1: A detailed distribution of unknown contigs across all microbiomes where each microbiome is represented by a subplot in the facet plot.

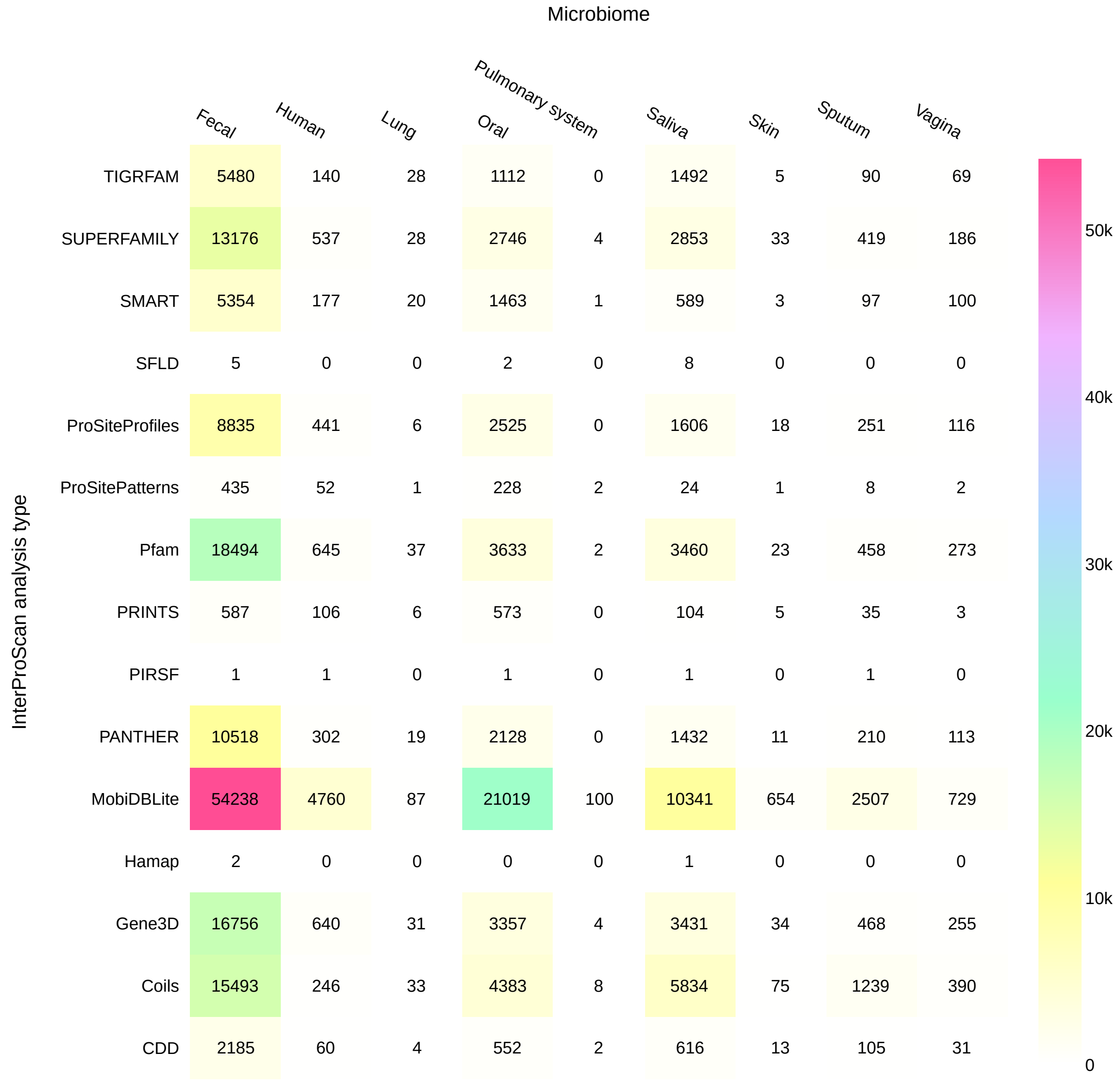


Figure S2: Functional homologues identified in the InterProScan analyses for UCs generated for each microbiome. Darker colours represent the higher number of hits to specific Pfam clans.

| Pfam clan description | Fecal | Saliva | Oral | Human | Sputum | Vagina |
|---|---|---|---|---|---|---|
| Tetratrico peptide repeat superfamily | 3316 | 261 | 462 | 98 | 35 | 25 |
| Leucine Rich Repeat | 2924 | 656 | 411 | | 34 | 40 |
| Choline binding repeat superfamily | 1154 | 340 | 288 | 80 | 23 | 27 |
| MORN repeat | 614 | 117 | 140 | | 51 | 41 |
| Helix-turn-helix clan | 489 | 74 | 91 | | | |
| Pectate lyase-like beta helix | 412 | | 53 | | | |
| Ig-like fold superfamily (E-set) | 405 | 49 | 49 | | | |
| Zinc beta-ribbon | 356 | | 23 | | | |
| Ankyrin repeat superfamily | 42 | | 294 | | | |
| Ubiquitin superfamily | 275 | 40 | 32 | | | |
| Beta propeller clan | 203 | | 36 | 36 | | |
| P-loop containing nucleoside triphosphate hydrolase superfamily | 179 | 165 | 78 | | | |
| beta-strand roll of R-module, superfamily | 142 |  | 179 | 91 |  |  |
| Pentapeptide repeat | 154 | | 38 | | | |
| EF-hand like superfamily | 141 | | | | | |
| Transthyretin superfamily | 128 | 21 | 26 | | | |
| Periplasmic binding protein clan | 125 | | | | | |
| Carbohydrate binding domain superfamily | 101 | | | | | |
| Src homology-3 domain | 95 | | | | | |
| PD-(D/E)XK nuclease superfamily | 91 | | 24 | | | |
| Galactose-binding domain-like superfamily | 75 | | | | | |
| Peptidase clan CA | 72 | 32 | 26 | | | |
| Glycosyl transferase GT-C superfamily | 72 | | | | | |
| Major Facilitator Superfamily | 60 | | | | | |
| Cupin fold | 54 | | | | | |
| Ribonuclease H-like superfamily | 44 | 23 | | | | |
| OB fold | 43 | | | | | |
| Concanavalin-like lectin/glucanase superfamily | 43 | | | | | |
| Tim barrel glycosyl hydrolase superfamily | 39 | | | | | |
| Fimbriae A and Mfa superfamily | 38 | | | | | |
| Barrel sandwich hybrid superfamily | 37 | | | | | |
| Hexapeptide repeat superfamily | 36 | | | | | |
| lambda integrase N-terminal domain | 36 | | | | | |
| Mirror Beta Grasp superfamily | 35 | | | | | |
| N-acetyltransferase like | 34 | | | | | |
| FAD/NAD(P)-binding Rossmann fold Superfamily | 33 | | | | | |
| Calcineurin-like phosphoesterase superfamily | 33 | | | | | |
| Peptidase clan MA | 33 | 24 | | | | |
| DNA polymerase B like | | 31 | | | | |
| PH domain-like superfamily | 29 | | | | | |
| O-antigen assembly enzyme superfamily | 29 | | | | | |
| Beta-trefoil superfamily | 28 | | | | | |
| Nucleotide cyclase superfamily | 28 | | | | | |
| Outer membrane beta-barrel protein superfamily | 28 | | | | | |
| ABC transporter membrane domain clan | 27 | | | | | |
| ABC-2-transporter-like clan | 24 | | | | | |
| Nucleotidyltransferase superfamily | 22 | | | | | |
| Rep-like domain | 22 | | | | | |
| Lysozyme-like superfamily | | 21 | | | | |
| Helix-hairpin-helix superfamily | 20 | | | | | |

Microbiome

Figure S3: Pfam clans found in different biomes

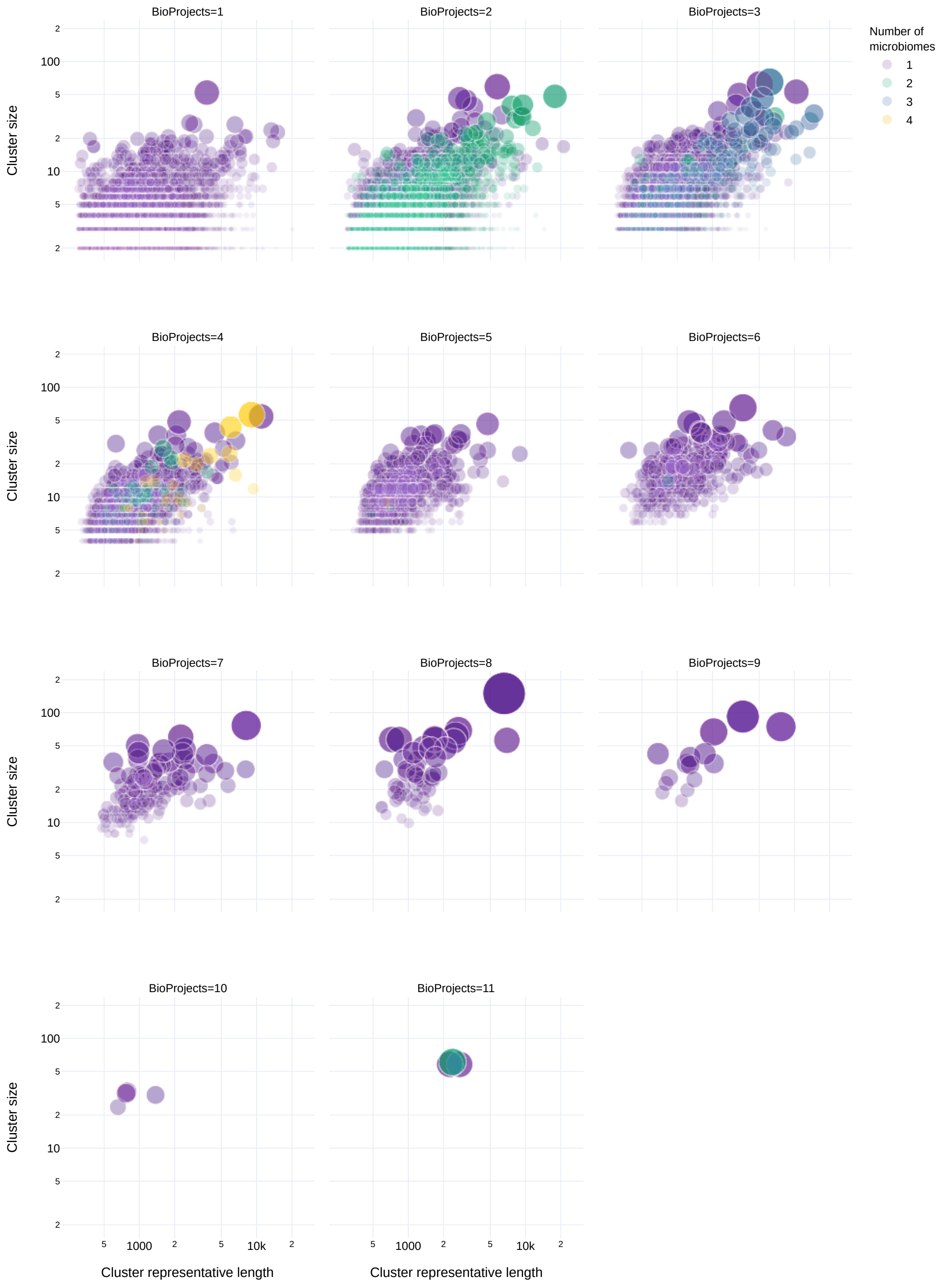


Figure S4: Overview of clusters found in the unknown sequence dataset. Number of biomes included in distinct clusters are represented by the columns and the number of bioprojects are represented by rows in the facetted plot. For each subplot, the X-axis represent the length of the cluster representative sequence and the Y-axis represent the cluster size for all cluster of size >=2.

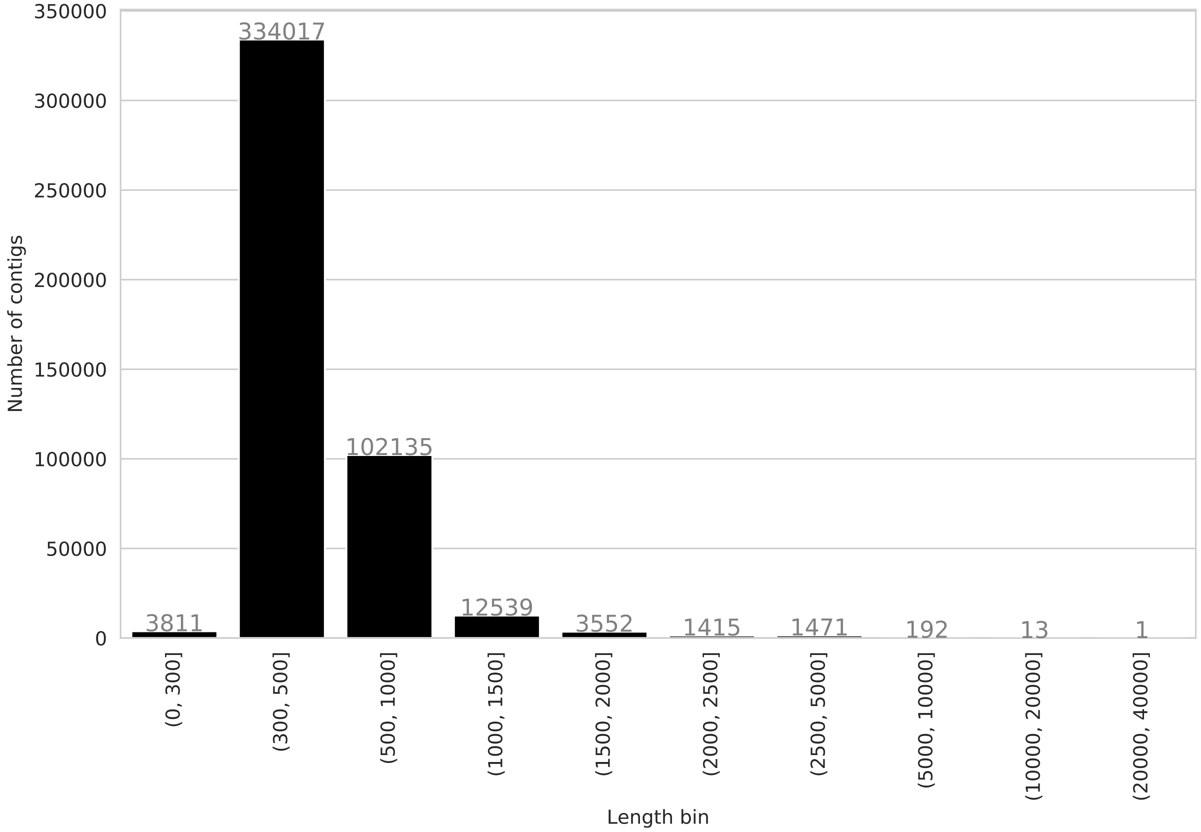


Figure S5: Distribution of contig lengths for all unknown contigs after the final time point (14 Oct 2020)

(a)

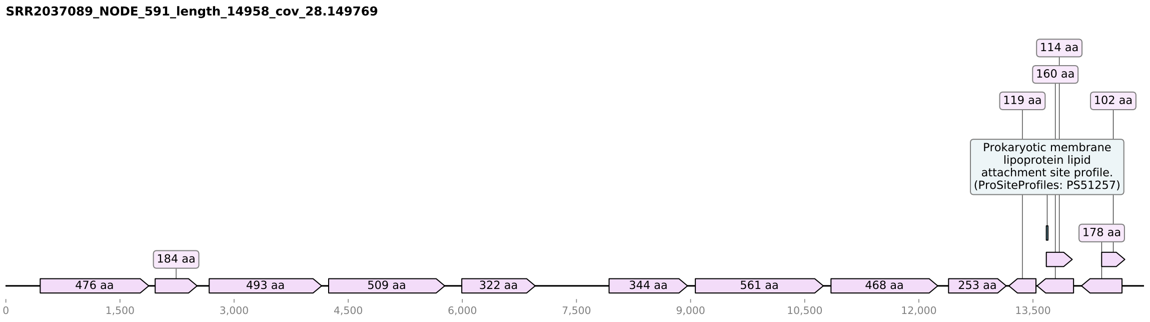


(b)

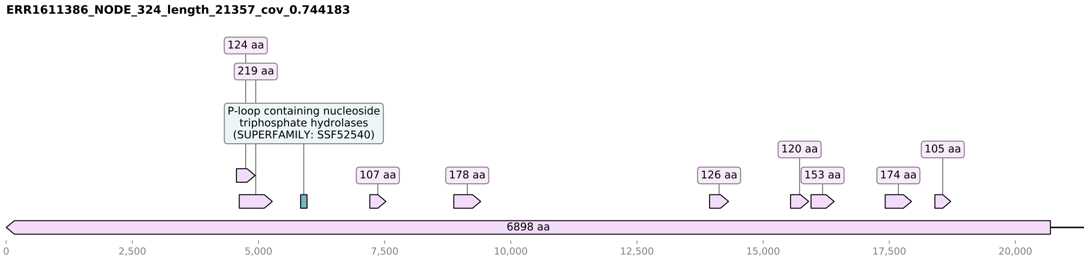


(c)

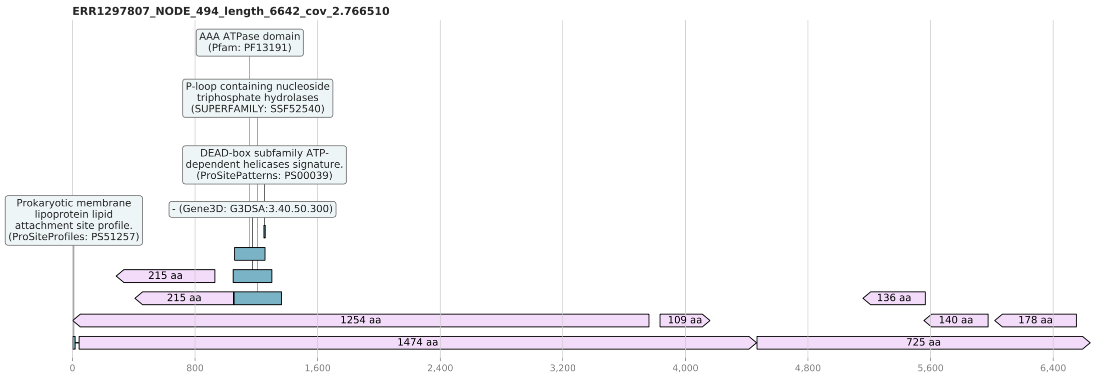


Figure S6: The genome diagrams of large unknown contigs (UCs) showing the open reading frames (ORFs) in light pink shade with the ORFs lengths as their corresponding labels and the green boxes illustrating the InterProScan predicted presence of domain signature. (a) Cluster representative of the largest cluster comprising UCs across multiple micobiomes. (b) UC with the largest predicted ORF (6,898 AA). (c) Cluster representative of the largest single microbiome cluster derived from fecal microbiome. This cluster comprised of 153 unknown contigs assembled from 46 samples across 8 different studies.

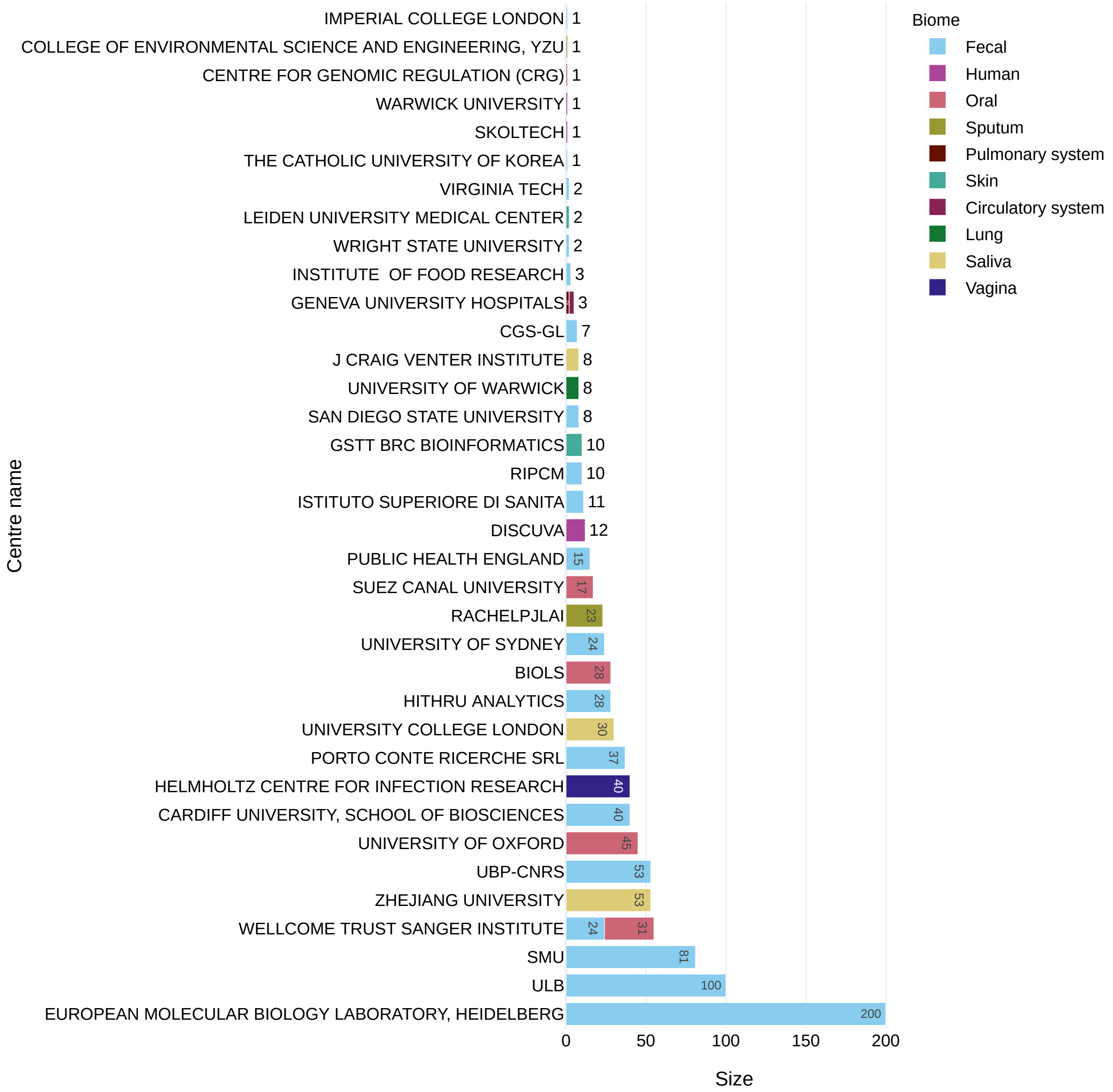


Figure S7: Research centres around the world coloured according to microbiome data sets distributed among them (n=963)

(a)

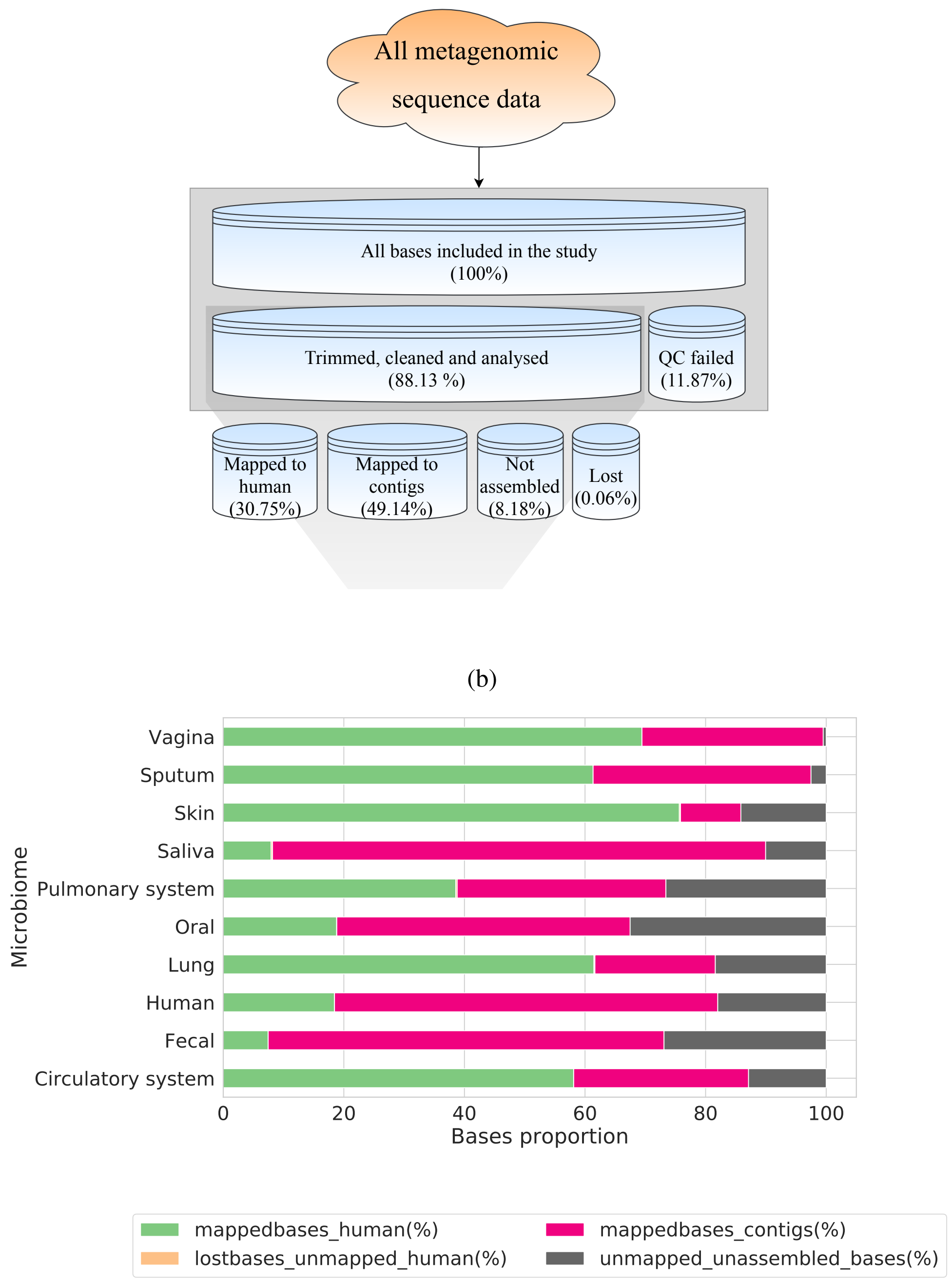


Figure S8: An overview of all bases analysed and categorised in this study. (a) Overall categorisation of all bases included in the study. (b) Proportion of bases mapped to human, assembled contigs and bases that were not assembled in different biomes.

### Supplementary tables

Table S1: List of samples excluded from the metagenomic analysis

| BioProject | Total number of samples | Samples excluded | Reason for exclusion |
|---|---|---|---|
| PRJEB14383 | 31 | 1 | raw sequence data unavailable (PE) |
| PRJEB14782 | 93 | 93 | not assembled (SE) |
| PRJEB15057 | 56 | 56 | not assembled (SE) |
| PRJEB15334 | 48 | 3 | not assembled (SE) |
| PRJEB19090 | 38 | 1 | not assembled (PE) |
| PRJEB19188 | 8 | 8 | raw sequence data unavailable (PE) |
| PRJEB20595 | 4 | 4 | not assembled (PE) |
| PRJEB21696 | 2 | 1 | not assembled (SE) |

Table S2: List of sample accession and the corresponding BioProject identifiers of samples excluded from the metagenomic analysis.

| BioProject | Run |
|---|---|
| PRJEB14782 | ERR1529686 |
| PRJEB14782 | ERR1529669 |
| PRJEB14782 | ERR1529668 |
| PRJEB14782 | ERR1529667 |
| PRJEB14782 | ERR1529666 |
| PRJEB14782 | ERR1529665 |
| PRJEB14782 | ERR1529664 |
| PRJEB14782 | ERR1529670 |
| PRJEB14782 | ERR1529663 |
| PRJEB14782 | ERR1529661 |
| PRJEB14782 | ERR1529660 |
| PRJEB14782 | ERR1529659 |
| PRJEB14782 | ERR1529658 |
| PRJEB14782 | ERR1529657 |
| PRJEB14782 | ERR1529656 |
| PRJEB14782 | ERR1529662 |
| PRJEB14782 | ERR1529671 |
| PRJEB14782 | ERR1529672 |
| PRJEB14782 | ERR1529673 |
| PRJEB14782 | ERR1529688 |
| PRJEB14782 | ERR1529687 |

| BioProject | Run |
|---|---|
| PRJEB14782 | ERR1529615 |
| PRJEB14782 | ERR1529685 |
| PRJEB14782 | ERR1529684 |
| PRJEB14782 | ERR1529683 |
| PRJEB14782 | ERR1529682 |
| PRJEB14782 | ERR1529681 |
| PRJEB14782 | ERR1529680 |
| PRJEB14782 | ERR1529679 |
| PRJEB14782 | ERR1529678 |
| PRJEB14782 | ERR1529677 |
| PRJEB14782 | ERR1529676 |
| PRJEB14782 | ERR1529675 |
| PRJEB14782 | ERR1529674 |
| PRJEB14782 | ERR1529655 |
| PRJEB14782 | ERR1529654 |
| PRJEB14782 | ERR1529653 |
| PRJEB14782 | ERR1529652 |
| PRJEB14782 | ERR1529632 |
| PRJEB14782 | ERR1529631 |
| PRJEB14782 | ERR1529630 |
| PRJEB14782 | ERR1529629 |
| PRJEB14782 | ERR1529628 |
| PRJEB14782 | ERR1529627 |
| PRJEB14782 | ERR1529626 |
| PRJEB14782 | ERR1529625 |
| PRJEB14782 | ERR1529624 |
| PRJEB14782 | ERR1529623 |
| PRJEB14782 | ERR1529622 |
| PRJEB14782 | ERR1529621 |
| PRJEB14782 | ERR1529620 |
| PRJEB14782 | ERR1529619 |
| PRJEB14782 | ERR1529618 |
| PRJEB14782 | ERR1529633 |
| PRJEB14782 | ERR1529689 |
| PRJEB14782 | ERR1529634 |
| PRJEB14782 | ERR1529636 |
| PRJEB14782 | ERR1529651 |
| PRJEB14782 | ERR1529650 |
| PRJEB14782 | ERR1529649 |

| BioProject | Run |
|---|---|
| PRJEB14782 | ERR1529648 |
| PRJEB14782 | ERR1529647 |
| PRJEB14782 | ERR1529646 |
| PRJEB14782 | ERR1529645 |
| PRJEB14782 | ERR1529644 |
| PRJEB14782 | ERR1529643 |
| PRJEB14782 | ERR1529642 |
| PRJEB14782 | ERR1529641 |
| PRJEB14782 | ERR1529640 |
| PRJEB14782 | ERR1529639 |
| PRJEB14782 | ERR1529638 |
| PRJEB14782 | ERR1529637 |
| PRJEB14782 | ERR1529635 |
| PRJEB14782 | ERR1529617 |
| PRJEB14782 | ERR1529690 |
| PRJEB14782 | ERR1529692 |
| PRJEB14782 | ERR1529613 |
| PRJEB14782 | ERR1529614 |
| PRJEB14782 | ERR1529691 |
| PRJEB14782 | ERR1529705 |
| PRJEB14782 | ERR1529704 |
| PRJEB14782 | ERR1529703 |
| PRJEB14782 | ERR1529702 |
| PRJEB14782 | ERR1529701 |
| PRJEB14782 | ERR1529616 |
| PRJEB14782 | ERR1529699 |
| PRJEB14782 | ERR1529698 |
| PRJEB14782 | ERR1529697 |
| PRJEB14782 | ERR1529696 |
| PRJEB14782 | ERR1529693 |
| PRJEB14782 | ERR1529694 |
| PRJEB14782 | ERR1529700 |
| PRJEB14782 | ERR1529695 |
| PRJEB15057 | ERR1558907 |
| PRJEB15057 | ERR1558906 |
| PRJEB15057 | ERR1558905 |
| PRJEB15057 | ERR1558904 |
| PRJEB15057 | ERR1558901 |
| PRJEB15057 | ERR1558902 |

| BioProject | Run |
|---|---|
| PRJEB15057 | ERR1558908 |
| PRJEB15057 | ERR1558900 |
| PRJEB15057 | ERR1558899 |
| PRJEB15057 | ERR1558903 |
| PRJEB15057 | ERR1558909 |
| PRJEB15057 | ERR1558912 |
| PRJEB15057 | ERR1558911 |
| PRJEB15057 | ERR1558898 |
| PRJEB15057 | ERR1558913 |
| PRJEB15057 | ERR1558914 |
| PRJEB15057 | ERR1558915 |
| PRJEB15057 | ERR1558916 |
| PRJEB15057 | ERR1558917 |
| PRJEB15057 | ERR1558918 |
| PRJEB15057 | ERR1558919 |
| PRJEB15057 | ERR1558920 |
| PRJEB15057 | ERR1558921 |
| PRJEB15057 | ERR1558922 |
| PRJEB15057 | ERR1558910 |
| PRJEB15057 | ERR1558897 |
| PRJEB15057 | ERR1558894 |
| PRJEB15057 | ERR1558895 |
| PRJEB15057 | ERR1558867 |
| PRJEB15057 | ERR1558868 |
| PRJEB15057 | ERR1558869 |
| PRJEB15057 | ERR1558870 |
| PRJEB15057 | ERR1558871 |
| PRJEB15057 | ERR1558872 |
| PRJEB15057 | ERR1558873 |
| PRJEB15057 | ERR1558874 |
| PRJEB15057 | ERR1558875 |
| PRJEB15057 | ERR1558876 |
| PRJEB15057 | ERR1558877 |
| PRJEB15057 | ERR1558878 |
| PRJEB15057 | ERR1558896 |
| PRJEB15057 | ERR1558879 |
| PRJEB15057 | ERR1558881 |
| PRJEB15057 | ERR1558882 |
| PRJEB15057 | ERR1558883 |

| **BioProject** | **Run** |
|---|---|
| PRJEB15057 | ERR1558884 |
| PRJEB15057 | ERR1558885 |
| PRJEB15057 | ERR1558886 |
| PRJEB15057 | ERR1558887 |
| PRJEB15057 | ERR1558888 |
| PRJEB15057 | ERR1558890 |
| PRJEB15057 | ERR1558891 |
| PRJEB15057 | ERR1558892 |
| PRJEB15057 | ERR1558893 |
| PRJEB15057 | ERR1558880 |
| PRJEB15057 | ERR1558889 |
| PRJEB15334 | ERR1611424 |
| PRJEB15334 | ERR1611401 |
| PRJEB15334 | ERR1611418 |
| PRJEB19090 | ERR1809127 |
| PRJEB20595 | ERR1951441 |
| PRJEB20595 | ERR1951440 |
| PRJEB20595 | ERR1951439 |
| PRJEB20595 | ERR1951438 |
| PRJEB21696 | ERR2028014 |

Table S3: Brief description of BioProjects included in the study.

| Biome | BioProject | Description | Library Layout | Samples | Reference |
|---|---|---|---|---|---|
| Circulatory system | PRJEB21816 | Aortite | PAIRED | 1 | Foulex et al., 2019 |
| Circulatory system | PRJEB24753 | Endocarditis due to Neisseria meningitidis | PAIRED | 2 | |
| Fecal | PRJEB10865 | Study of the abundance of bacteria from human samples | PAIRED | 2 | |
| Fecal | PRJEB11554 | NeoM | PAIRED | 1 | |
| Fecal | PRJEB12357 | Impact of faecal microbiota transplantation on the intestinal microbiome in metabolic syndrome patients | PAIRED | 100 | Li et al., 2016 |
| Fecal | PRJEB14935 | Term and preterm shotgun samples | PAIRED | 3 | Alcon-Giner et al., 2017 |
| Fecal | PRJEB15257 | The antibiotic resistance potential of the preterm infant gut microbiome measured using shotgun metagenomics. | PAIRED | 15 | Rose et al., 2017 |
| Fecal | PRJEB1775 | Diagnostic Metagenomics: A Culture-Independent Approach to the Investigation of Bacterial Infections | PAIRED | 53 | Loman et al., 2013 |
| Fecal | PRJEB17784 | The fecal microbiota in L-DOPA naive PD patients | SINGLE | 100 | |
| Fecal | PRJEB18265 | Estimation of variability in the gut microbiota resistome of the Russian citizens aimed at identification of pathways for transmission and spread of antibiotic resistance. | PAIRED | 10 | Olekhnovich et al., 2019 |
| Fecal | PRJEB19090 | Potential and active functions in the gut microbiota of a healthy human cohort | PAIRED | 37 | Tanca et al., 2017 |

| Biome | BioProject | Description | Library Layout | Samples | Reference |
|---|---|---|---|---|---|
| Fecal | PRJEB19367 | Analysis of stool samples from sickle cell disease patients and healthy controls | PAIRED | 28 | |
| Fecal | PRJEB21696 | Metagenomics 1st 5 data | SINGLE | 1 | |
| Fecal | PRJEB23207 | Metagenomic characterization of the human intestinal microbiota in faecal samples from STEC-infected patients | PAIRED | 11 | |
| Fecal | PRJEB5761 | Gut microbiota in chronic kidney disease | PAIRED | 81 | |
| Fecal | PRJEB6092 | Metagenome fecal microbiota- Illumina seq reads of 12 individuals at 2 timepoints | PAIRED | 24 | |
| Fecal | PRJEB6542 | Gut microbial metabolism shifts towards a more toxic profile with supplementary iron in a kinetic model of the human large intestine | PAIRED | 8 | |
| Fecal | PRJEB7331 | metagenomic analysis of human gut microbiome | PAIRED | 24 | |
| Fecal | PRJEB7949 | The fecal microbiome was studied in a group of IBD suffers and compared to a control group's fecal microbiome | PAIRED | 40 | |
| Fecal | PRJEB8094 | The initial state of the human gut microbiome determines its reshaping by antibiotics | PAIRED | 100 | Raymond et al., 2016 |
| Fecal | PRJEB8201 | Comparison of distal gut microbiota structure and function in US and Egyptian children | SINGLE | 2 | |
| Fecal | PRJNA43253 | Human fecal microbiome | SINGLE | 7 | Turnbaugh et al., 2010 |
| Human | PRJEB14301 | CSF | SINGLE | 1 | |
| Human | PRJEB21827 | A/B testing for colon model | PAIRED | 12 | |

| Biome | BioProject | Description | Library Layout | Samples | Reference |
|---|---|---|---|---|---|
| Human | PRJEB6045 | metagenomics of medieval human remains from Sardinaia | PAIRED | 1 | |
| Lung | PRJEB7248 | Metagenomics of TB-associated sputum | PAIRED | 8 | |
| Oral | PRJEB12831 | A plaque on both your houses. Exploring the history of urbanisation and infectious diseases through the study of archaeological dental tartar | PAIRED | 31 | |
| Oral | PRJEB12998 | Test file for Oralfungi project | PAIRED | 1 | |
| Oral | PRJEB15334 | Radcliffe dental calculus | SINGLE | 45 | |
| Oral | PRJNA230363 | Oral Microbiome | PAIRED | 28 | |
| Oral | PRJNA384402 | oral metagenome Metagenome | PAIRED | 17 | |
| Pulmonary system | PRJEB20877 | Detection of bacterial pathogens from broncho-alveolar lavage by next-generation sequencing | PAIRED | 2 | Leo et al., 2017 |
| Saliva | PRJEB14383 | Oral microbiome samples from the Philippines | PAIRED | 30 | Lassalle et al., 2018 |
| Saliva | PRJNA264728 | Gene expression anlayses of saliva-derived in vitro biofilms during carbohydrate fermentation and pH stress | PAIRED | 8 | Edlund et al., 2015 |
| Saliva | PRJNA306560 | Human oral saliva Metagenome | PAIRED | 53 | |
| Skin | PRJEB10133 | These samples are selections from a larger cohort that were selected for the participation in the EBI metagenomics training in Sept. 2015 | PAIRED | 10 | |
| Skin | PRJEB10295 | Whole genome sequencing of metagenomes extracted from palms of two individuals | PAIRED | 2 | |
| Sputum | PRJEB10919 | Total RNA-Seq on sputum samples from patients with active tuberculosis | SINGLE | 23 | Schnettger et al., 2017 |

| Biome | BioProject | Description | Library Layout | Samples | Reference |
|---|---|---|---|---|---|
| Sputum | PRJEB14539 | Metagenome and metatranscriptome profiling of moderate and severe COPD sputum in Taiwanese Han males | PAIRED | 1 | |
| Vagina | PRJEB21446 | Metatranscriptome reveals the function shifts of the vaginal microbiome during the treatment of bacterial vaginosis | PAIRED | 40 | Deng et al., 2018 |

Table S4: List of studies with the country of origin and biome details .

| Country | BioProject | Biome | Count |
|---|---|---|---|
| Australia | PRJEB6092 | Fecal | 24 |
| Canada | PRJEB8094 | Fecal | 100 |
| China | PRJNA230363 | Oral | 28 |
| China: Sichuan | PRJNA306560 | Saliva | 53 |
| Egypt | PRJEB8201 | Fecal | 1 |
| Egypt | PRJNA384402 | Oral | 17 |
| Gambia | PRJEB7248 | Lung | 8 |
| Germany | PRJEB1775 | Fecal | 53 |
| Germany | PRJEB17784 | Fecal | 100 |
| Germany | PRJEB21446 | Vagina | 40 |
| Italy | PRJEB19090 | Fecal | 37 |
| Italy | PRJEB23207 | Fecal | 7 |
| Italy: Sardinia | PRJEB6045 | Human | 1 |
| Netherlands | PRJEB10295 | Skin | 2 |
| Netherlands | PRJEB12357 | Fecal | 100 |
| Netherlands | PRJEB23207 | Fecal | 4 |
| Netherlands | PRJEB6542 | Fecal | 8 |
| Philippines | PRJEB14383 | Saliva | 30 |
| Russia | PRJEB14301 | Human | 1 |
| Russia | PRJEB18265 | Fecal | 10 |
| South Africa | PRJEB10919 | Sputum | 19 |
| South Korea | PRJEB21696 | Fecal | 1 |
| Switzerland | PRJEB20877 | Pulmonary system | 2 |
| Switzerland | PRJEB21816 | Circulatory system | 1 |
| Switzerland | PRJEB24753 | Circulatory system | 2 |
| Taiwan | PRJEB14539 | Sputum | 1 |
| USA | PRJEB10865 | Fecal | 2 |
| USA | PRJEB19367 | Fecal | 28 |
| USA | PRJEB8201 | Fecal | 1 |
| United Kingdom | PRJEB10133 | Skin | 10 |
| United Kingdom | PRJEB10919 | Sputum | 4 |
| United Kingdom | PRJEB12831 | Oral | 31 |
| United Kingdom | PRJEB14935 | Fecal | 3 |
| United Kingdom | PRJEB15257 | Fecal | 15 |
| United Kingdom | PRJEB15334 | Oral | 45 |
| United Kingdom | PRJEB7331 | Fecal | 24 |
| United Kingdom | PRJEB7949 | Fecal | 40 |

| Country | BioProject | Biome | Count |
|---|---|---|---|
| nan | PRJEB11554 | Fecal | 1 |
| nan | PRJEB12998 | Oral | 1 |
| nan | PRJEB21827 | Human | 12 |
| nan | PRJEB5761 | Fecal | 81 |
| nan | PRJNA43253 | Fecal | 7 |
| not applicable | PRJNA264728 | Saliva | 8 |

Table S5: Average read lengths of raw data vs assembled sequences .

| | Average read length(SRA) | Average read length (Contigs) |
|---|---|---|
| count | 963.0 | 963.0 |
| mean | 129.0 | 108.89 |
| std | 58.57 | 40.88 |
| min | 36.0 | 30.0 |
| max | 301.0 | 264.0 |

Table S6: Proportion of bases lost after QC .

| | Circulatory system | Fecal | Human | Lung | Oral | Pulmonary system | Saliva | Skin | Sputum | Vagina |
|---|---|---|---|---|---|---|---|---|---|---|
| count | 3.0 | 647.0 | 14.0 | 8.0 | 122.0 | 2.0 | 91.0 | 12.0 | 24.0 | 40.0 |
| mean | 5.94 | 10.48 | 20.11 | 23.45 | 25.41 | 21.01 | 27.66 | 15.26 | 7.72 | 1.95 |
| std | 8.48 | 9.49 | 5.34 | 6.17 | 17.36 | 20.78 | 23.37 | 12.54 | 2.24 | 0.23 |
| min | 0.86 | 0.03 | 15.1 | 13.99 | 2.78 | 6.32 | 0.8 | 5.19 | 3.02 | 1.64 |
| max | 15.73 | 63.94 | 34.8 | 33.44 | 94.63 | 35.7 | 77.03 | 50.83 | 13.26 | 2.55 |
